# Temporoparietal alpha dynamics shape feedback-based updating during social bargaining

**DOI:** 10.64898/2026.04.03.716401

**Authors:** Alejandra Figueroa-Vargas, Gabriela Valdebenito-Oyarzo, Maria Paz Martinez-Molina, Patricia Soto-Icaza, Paulo Figueroa-Taiba, Marcela Díaz-Díaz, Matias Iriarte-Carter, César Salinas, Ximena Stecher, Carla Manterola, Sebastián Navarrete, Pamela Guevara, Francisco Zamorano, Antoni Valero-Cabre, Rafael Polania, Pablo Billeke

## Abstract

Adaptive behavior requires weighting incoming feedback against more stable action policies, but the neural dynamics controlling this balance during social interaction remain unclear. Across 296 experimental sessions in 219 participants, we combined computational modeling, EEG, fMRI, diffusion MRI, and fMRI-guided rhythmic TMS during repeated social bargaining. Behavior was captured by complementary feedback-based updating and reputation-based adjustment, with reproducible individual differences in how instructed social context reweighted feedback. Feedback-based updating showed the clearest convergent neural signature: trial-by-trial updating was associated with alpha–beta dynamics and multivoxel representational structure in right temporoparietal cortex. Critically, rhythmic alpha-TMS over individually localized TPJ sites selectively reduced feedback-based updating relative to active arrhythmic stimulation, while increasing baseline offers and modulating anticipatory alpha activity. Functional and structural connectivity analyses further implicated a distributed temporoparietal–frontal network. Together, these findings link a specific computational operation—feedback weighting—to temporoparietal alpha dynamics and show that rhythmic perturbation alters its behavioral expression.

## Introduction

Successful negotiation requires individuals to anticipate others’ responses, update expectations from previous interactions, and adapt their behavior accordingly. These processes operate across contexts ranging from everyday interpersonal exchanges to group and political negotiations. Yet the computational and neural mechanisms supporting such adaptive social learning remain incompletely understood.

Strategic social interactions require agents to update predictions about others in environments where outcomes depend on both their own and others’ actions. Repeated strategic games provide a controlled and computationally tractable framework for studying this process, requiring individuals to continuously update beliefs about an interaction partner and use them to guide future actions ^1–6^. Importantly, adaptive social behavior does not emerge from a uniform strategy across individuals ^7^. People display consistent differences in how they use social information and adopt learning strategies ^8^, suggesting that population-level averages may obscure meaningful variation in how feedback guides behavior. Computational accounts similarly emphasize individual differences in the weighting of new feedback, prior expectations, and beliefs about others’ changing behavior^9^. Repeated bargaining therefore provides a useful framework for examining both how people adapt to social feedback and how different adaptive strategies vary across individuals.

These computations rely on distributed fronto–temporoparietal networks^5,10^. Prefrontal regions have been implicated in value integration, model-based control, and strategic planning^6,11^, whereas the temporoparietal junction (TPJ) contributes to tracking others’ behavior and updating beliefs about action–outcome contingencies and others’ responsiveness ^5,6,12,13^. Prior work using repeated ultimatum games has identified that alpha and beta oscillations in the right temporoparietal cortex are associated with anticipating others’ responses during negotiation and correlate with subsequent player behavior ^14,15^. These dynamics interact with frontal activity, mainly with theta activity in relation to the evaluation of others’ responses ^16^ in accordance with the role of this frequency in learning and cognitive control expectations ^11,17,18^. Beyond alpha-band activity, frequency-specific temporoparietal mechanisms have been implicated in perspective-taking, inequity aversion, and prosocial behavior through theta- and gamma-band dynamics^19,20^. Together, this evidence suggests that adaptive social behavior depends on frequency-specific dynamics within temporoparietal–frontal networks. However, most evidence derives from isolated judgments or one-shot decisions, leaving unresolved how these dynamics support more specifically feedback-based adaptation across repeated interactions and whether experimentally perturbing them alters learning.

Based on evidence that individuals differ in their use of social learning strategies^7–9^, together with previous correlational findings implicating the TPJ and alpha oscillations in strategic social interaction^14,21,22^, we asked whether feedback-based updating could be isolated computationally, mapped onto temporoparietal oscillatory dynamics, and selectively perturbed. We hypothesized that feedback-based updating would be particularly associated with alpha-band dynamics within a TPJ-centered network and that rhythmic perturbation of these dynamics would alter the behavioral weight assigned to social feedback. To test this framework, we combined computational modeling across independent samples with EEG, fMRI representational similarity analysis, and fMRI-guided rhythmic TMS-EEG.

## Results

### Experimental rationale

We organized the experiments and analyses according to the sequential logic of the research program (Figure 1B). The initial EEG sample was used to characterize the computational strategies underlying bargaining behavior and their oscillatory signatures, including individual differences in strategy modulation across social contexts. Independent behavioral and Human-only samples then tested the generalization of this computational architecture. The fMRI sample provided anatomical and representational evidence for the neural implementation of feedback-based updating, while the TMS-EEG experiment tested whether rhythmic alpha stimulation of the TPJ-centered network could alter the corresponding computational and neural signatures. Functional and structural connectivity analyses further assessed whether these effects were expressed at the network level.

**Figure 1.**
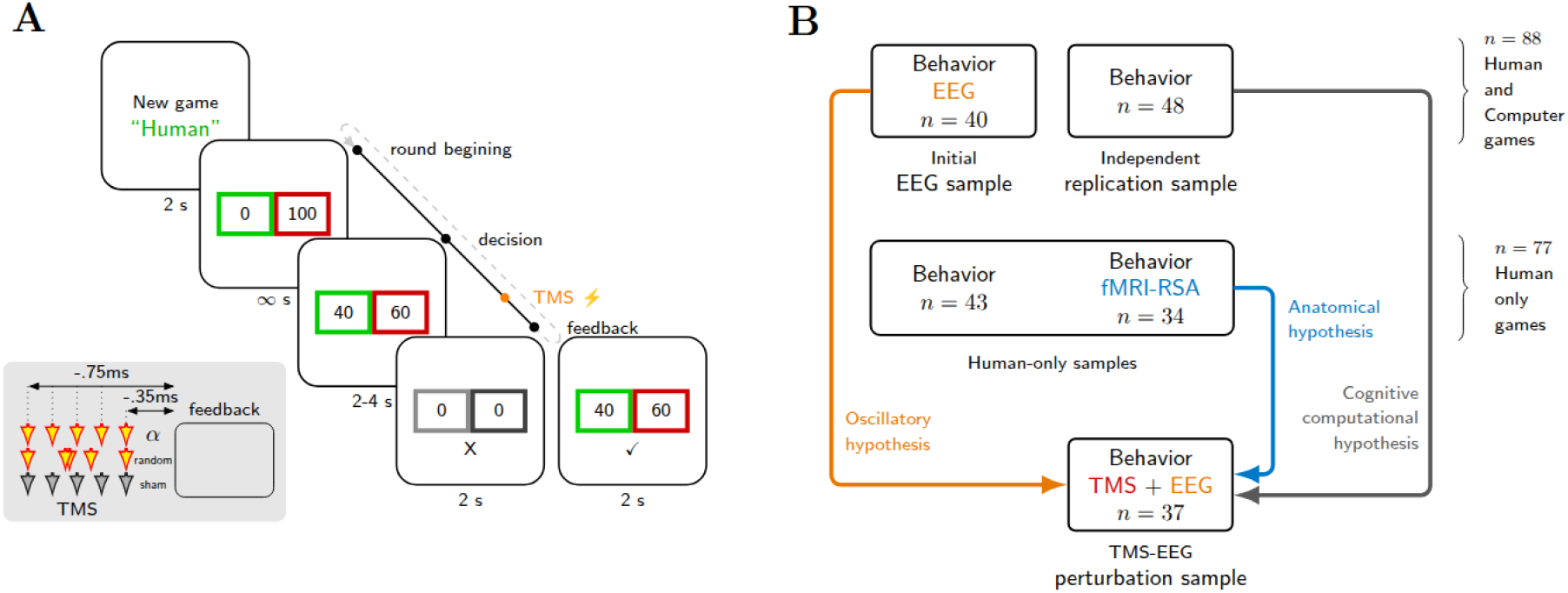
Experimental design and multimodal framework. **A,** Trial structure of the Ultimatum Game. Each round began with the presentation of an opponent (“Human” or “Computer” responder), followed by the decision phase in which participants selected an offer. A burst of five TMS pulses was delivered during the decision–feedback interval in the TMS experiment. Feedback indicating acceptance or rejection by the responder was then presented. The timing of each event is indicated. **B,** Overview of the experimental framework across samples. An initial EEG sample (n = 40) was used to characterize the computational strategies underlying bargaining behavior, individual differences in their modulation across social contexts, and their oscillatory signatures. An independent replication sample (n = 48) and Human-only samples (n = 77; 43 behavioral and 34 fMRI) tested the generalization of this computational architecture, with the fMRI sample additionally assessing the anatomical and representational correlates of feedback-based updating. Finally, a TMS-EEG sample (n = 37) tested whether rhythmic alpha stimulation of individually localized TPJ sites perturbed the corresponding computational and neural signatures.

### Behavior

We first conducted an EEG experiment with 40 participants aged 17–26 during a repeated Ultimatum Game (Figure 1A). Participants always acted as proposers and completed eight 20-round games with the same anonymous responder within each game. They played under two instructed contexts: in the Human condition, they were told that the responder was another participant interacting online, whereas in the Computer condition they were told that responses were generated by a probabilistic algorithm whose acceptance probability increased with offer size. In both conditions, responses were algorithmically generated using contingencies derived from previous human–human interaction studies^14^, allowing behavioral contingencies to be matched while manipulating the instructed identity of the responder.

In prior work using the same experimental setting, we showed that participants progressively adapted their offers across rounds to identify proposals that maximized both acceptance probability and earnings^21^. Here, we used the programmed acceptance probability associated with each offer as a proxy for behavioral adaptation during bargaining. As an initial descriptive analysis (Model M0), we examined changes in this measure across rounds and whether these differed between the Human and Computer conditions.

Hierarchical Bayesian analysis revealed that participants significantly increased their adaptation across game rounds (β_round_ median = 0.072; 95% High-Density Interval [HDI_95%_] = [0.028, 0.11]; p_MCMC_ = 0.002). At the population level, this change did not differ significantly between Human and Computer conditions (β_round,h_ = –0.029; HDI_95%_ = [–0.08, 0.015]; p_MCMC_ = 0.17). However, including Human-versus-Computer modulation substantially improved model fit and predictive accuracy (ΔDIC = −168; ΔLOOIC = −172), motivating subsequent analyses of whether social context was associated with structured individual differences in adaptive strategy use.

### Computational Strategy Models

To investigate the computational processes underlying adaptive behavior during repeated bargaining, we compared candidate models implementing two complementary strategic mechanisms derived from previous work on repeated games and social bargaining^12,14,16,23^. The first, termed the U-strategy, captured feedback-based updating: participants adjusted their behavior according to the accumulated history of acceptance and rejection outcomes. The second, termed the A-strategy, captured reputation-based adjustment: participants’ current behavior was influenced by the history of their own previous offers, providing a proxy for strategic adaptation aimed at shaping the responder’s future behavior. Full model specifications and learning rules are provided in the Methods section.

We asked whether these strategies contributed independently to bargaining behavior and whether their influence differed according to instructed responder identity. Candidate models therefore included the U- and A-strategies either separately or jointly, with or without Human-versus-Computer modulation.

### Model Comparison

To determine which computational processes best explained participants’ behavior, we compared candidate models incorporating the U- and A-strategies, either alone or in combination, and allowing their contributions to vary with instructed responder identity. The model combining feedback-based updating (U-strategy) and reputation-based adjustment (A-strategy), together with their Human-versus-Computer modulation, provided the best fit and predictive accuracy, as indicated by the lowest DIC and LOOIC values.

In this model, the U-strategy showed a significant population-level effect, indicating that feedback-based updating contributed to behavior across games (β_U_ median = 1.02; 95% HDI_95%_ = [0.2, 1.7]; p_MCMC_ = 0.012). The average modulation of this strategy by instructed responder identity was not significant (β_U,h_ median = 0.2; 95% HDI_95%_ = [–0.6, 1.2]; p_MCMC_ = 0.6) Similarly, the A-strategy contributed significantly to behavior across games (β_A_ median = 0.37; 95% HDI_95%_ = [0.24, 0.51]; p_MCMC_ < 0.001), whereas its average Human-versus-Computer modulation was close to zero (β_A,h_ median = −0.1; 95% HDI_95%_ = [–0.2, 0.05]; p = 0.19).

To assess whether the improved predictive performance associated with responder identity reflected meaningful condition structure rather than additional model flexibility, we repeated the analysis after permuting Human and Computer labels across games within participants. The observed model yielded a lower LOOIC than models fitted to permuted condition labels (permutation p = 0.0002), indicating that instructed responder identity contained information relevant for predicting participants’ behavior despite the absence of a homogeneous population-level shift in strategy coefficients. Because the hierarchical model allowed Human-versus-Computer modulation to vary across participants, opposing or differently sized individual effects could average to a group-level coefficient near zero. Thus, the absence of a population-level interaction should not be interpreted as evidence that responder identity did not influence strategy use. Instead, the superior predictive performance of the model preserving the true condition structure motivated us to examine these individual differences directly. We therefore tested whether participant-specific changes in computational strategy were systematically related to changes in bargaining behavior and whether distinct patterns of strategy modulation could be identified across participants. All model parameters were reliably recovered from simulated data, supporting the identifiability of the latent variables (Supplementary Figure 1), and model fits to the empirical data are shown in Supplementary Figure 2.

**Table 1:** Summary of the models parameters.

| Model | Free parameters |
| --- | --- |
| M0 | **4**<br>$\beta_0$ (intercept), $\beta_r$ (rounds), $\beta_h$ (human games), $\beta_{r,h}$ (interaction round and human games ) |
| Muh | **6**<br>$\beta_0$ (intercept), $\beta_u$ (U-strategy), $\alpha_{Ur}$ (Learning rate for rejection), $\alpha_{Ua}$ (Learning rate for acceptance), $\beta_h$ (human games), and $\beta_{U,hum}$ (interaction U-strategy and human games ) |
| Mah | **6**<br>$\beta_0$ (intercept), $\beta_A$ (A-strategy), $\alpha_{Ar}$ (Learning rate for reputation following rejection), $\alpha_{Aa}$ (Learning rate for reputation following acceptance), $\beta_h$ (human games), and $\beta_{A,hum}$ (interaction A-strategy and human games ). |
| Mua | **7**<br>$\beta_0$ (intercept), $\beta_u$ (U-strategy), $\alpha_{Ur}$ (Learning rate for rejection), $\alpha_{Ua}$ (Learning rate for acceptance), $\beta_A$ (A-strategy), $\alpha_{Ar}$ (Learning rate for reputation following rejection) and $\alpha_{Aa}$ (Learning rate for reputation following acceptance). |
| Mua_ | **8**<br>$\beta_0$ (intercept), $\beta_u$ (U-strategy), $\alpha_{Ur}$ (Learning rate for rejection), $\alpha_{Ua}$ (Learning rate for acceptance), |
| M0 | **4**<br>$\beta_0$ (intercept), $\beta_r$ (rounds), $\beta_h$ (human games), $\beta_{r,h}$ (interaction round and human games ) |
| | $\beta_A$ (A-strategy), $\alpha_{Ar}$ (Learning rate for reputation following rejection), $\alpha_{Aa}$ (Learning rate for reputation following acceptance), and $\beta_h$ (human games). |
| Murh | **9**<br>$\beta_0$ (intercept), $\beta_u$ (U-strategy), $\alpha_{Ur}$ (Learning rate for rejection), $\alpha_{Ua}$ (Learning rate for acceptance), $\beta_R$ (R-strategy), $\alpha_R$ (Learning rate for reputation), $\beta_h$ (human games), $\beta_{U,hum}$ (interaction U-strategy and human games ), and $\beta_{A,hum}$ (interaction A-strategy and human games ). |
| Muah | **10**<br>$\beta_0$ (intercept), $\beta_u$ (U-strategy), $\alpha_{Ur}$ (Learning rate for rejection), $\alpha_{Ua}$ (Learning rate for acceptance), $\beta_A$ (A-strategy), $\alpha_{Ar}$ (Learning rate for reputation following rejection), $\alpha_{Aa}$ (Learning rate for reputation following acceptance), $\beta_h$ (human games), $\beta_{U,hum}$ (interaction U-strategy and human games ), and $\beta_{A,hum}$ (interaction A-strategy and human games ). |

### Individual differences in strategy modulation during Human games

The absence of population-level Human-versus-Computer interactions, together with the greater predictive performance obtained when preserving the true responder-identity structure, raised the possibility that social context affected strategy use differently across individuals. We therefore examined participant-specific modulation of the computational parameters during instructed Human games.

The clearest individual-level relationship emerged between the modulation of baseline offer level β_0,hum_ and feedback-based updating β_U,hum_. Participants who increased their baseline offers in Human games tended to show a stronger reduction in their reliance on feedback-driven updating (Spearman’s ρ = −0.82, p = 6x10^-11^). In contrast, baseline-offer modulation was not associated with modulation of the reputation-based A-strategy (Spearman’s ρ = 0.07, p = 0.6), and we found no evidence for a systematic association between Human-related changes in the U- and A-strategies (Spearman’s ρ = 0.2, p = 0.10). Thus, the strongest structure in the individual differences was a coordinated change in baseline bargaining behavior and sensitivity to responder feedback: participants who began Human games with higher offers subsequently relied less on feedback to adjust those offers, a pattern consistent with previous findings in repeated social bargaining ^16^.

We next examined whether these individual differences were associated with negotiation outcomes. Baseline offer level was positively associated with total earnings (β_0_: ρ = 0.54, p = 0.0002), and so was its modulation during Human games (β_0,hum_: ρ = 0.39, p = 0.01). Other strategy parameters showed no consistent associations with earnings (|ρ| < 0.28, p > 0.09). These findings indicate that individual variation in baseline bargaining behavior was behaviorally consequential, while its close relationship with feedback-based updating suggested a coordinated pattern of adaptation during instructed Human interactions.

### Exploratory analysis: latent heterogeneity in strategic adaptation

Given the structured individual differences observed in Human-versus-Computer modulation, we next asked whether the weak population-level interaction could arise because strategy changes were expressed more strongly in some participants than in others. We therefore fitted a hierarchical Bayesian latent-mixture model in which a participant-level latent variable allowed Human-related modulation of the computational parameters for individuals showing evidence of strategy change, while constraining this modulation toward zero for the remaining participants (see Methods).

The model estimated that approximately 79% of participants were assigned to the strategy-modulation component (SC median = 0.79, 95% HDI = [0.76, 0.80]). Within this component, instructed Human interactions were characterized by a selective reduction in feedback-based updating (β_U,Hum_ = −0.37, 95% HDI [−0.73, −0.09], p_MCMC_ = 0.007, Figure 2D) together with an increase in baseline offer level (β_0,Hum_ = 0.51, 95% HDI [0.18, 0.86], p_MCMC_ = 0.002, Figure 2D). In contrast, there was no evidence for systematic modulation of the reputation-based A-strategy (β_A,Hum_ = −0.07, 95% HDI [−0.50, 0.39], p_MCMC_ = 0.70).

**Figure 2.**
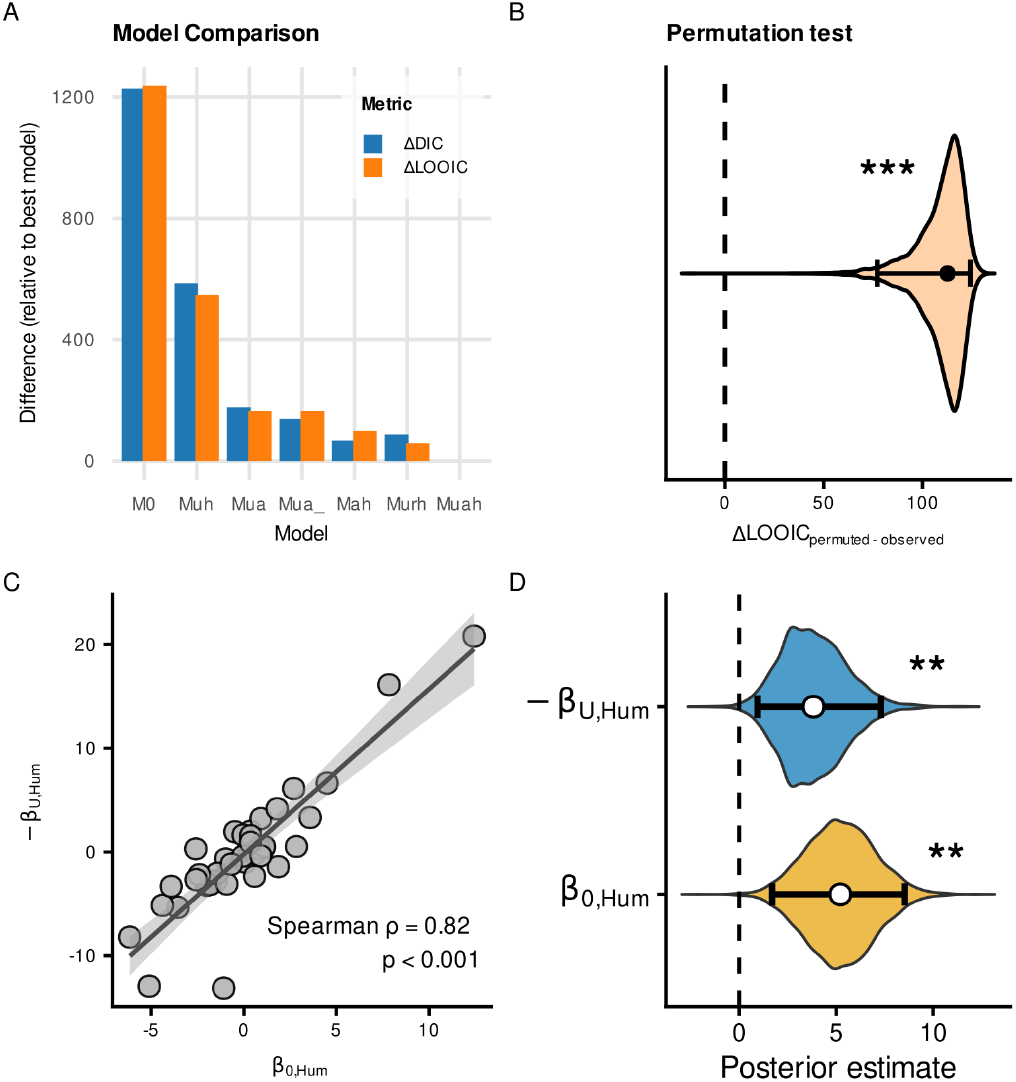
Computational and individual differences in behavioral strategies during social negotiation. **A,** Model comparison across candidate computational models. Bars show differences in model fit and predictive accuracy relative to the best-fitting model, quantified using ΔDIC and ΔLOOIC, respectively. The model combining feedback-based updating (U-strategy) and reputation-based adjustment (A-strategy), together with their modulation during instructed Human interactions, provided the best overall account of behavior. **B,** Permutation test assessing whether the predictive advantage associated with responder identity reflected meaningful Human-versus-Computer condition structure. The violin shows the distribution of ΔLOOIC values obtained after permuting condition labels across games within participants, expressed relative to the model fitted using the observed condition labels (permuted − observed). Positive values indicate poorer predictive performance after permutation. The dashed line indicates ΔLOOIC = 0. **C,** Individual differences in strategy modulation during Human games. Participants who increased their baseline offers more strongly during Human interactions (β_0,hum_) also showed a greater reduction in feedback-based updating, represented in the figure as −β_U,hum_. Points represent individual participants; the line and shaded area show the linear fit and 95% confidence interval for visualization. Spearman’s correlation is reported in the panel. **D,** Posterior distributions from the hierarchical Bayesian latent-mixture model for the Human-related increase in baseline offers (β_0,hum_) and reduction in feedback-based updating (β_U,hum_). White circles indicate posterior means and horizontal bars indicate 95% highest-density intervals (HDIs); the dashed vertical line indicates zero. Asterisks indicate posterior evidence against zero (*p_MCMC_<0.05, **p_MCMC_<0.01, ***p_MCMC_<0.001).

Importantly, the latent-mixture model provided a better account of participants’ behavior than the corresponding model assuming a common population-level modulation, improving both model fit and predictive accuracy (ΔDIC = −51; ΔLOOIC = −47). Thus, the weak average Human-versus-Computer effects were better explained by structured inter-individual variability than by a uniform shift in computational strategy. Specifically, a substantial subset of participants increased their baseline offers while simultaneously reducing their reliance on feedback-based updating during instructed Human interactions.

### Replication Samples

To determine whether these findings were specific to our initial sample or generalized across a broader population, we replicated the same analyses in an extended cohort of participants spanning a wide age range (11–30 years, *n* = 48) who completed the same experimental paradigm. This allowed us to assess the robustness and behavioral consistency of the observed effects across developmental stages, and to reduce the possibility that the computational architecture identified in the initial sample reflected sample-specific overfitting.

The main computational and individual-difference findings were replicated, although the relationship between the U- and A-strategy modulations differed between samples. The M_uah_ model provided the best fit and highest predictive accuracy (**Supplementary Figure 3**). The coefficients associated with human interactions (β_hum_) were not significant alone; however, preserving the true Human-versus-Computer condition structure significantly improved model predictive performance relative to permuted condition labels (permutation p <0.0001). Similarly, the latent-mixture model estimated that 83% of participants were assigned to the strategy-modulation component (SC, median = 0.83, 95% HDI [0.79, 0.88]) characterized by a decrease in U-strategy (β_U,Hum_ = −0.29, 95% HDI [−0.56, −0.03], p_MCMC_ = 0.027) and an increase in baseline offers during Human games (β_0,Hum_ = 0.4, 95% HDI [0.12, 0.69], p_MCMC_ = 0.003). No systematic shift was observed for the A-strategy (β_A,Hum_ = −0.26, 95% HDI [−0.67, 0.09], p_MCMC_ = 0.14). Moreover, in the replication sample, β_0,hum_ was significantly associated with β_U,hum_ (Spearman’s ρ = −0.54, p = 6 × 10−5), but not with β_A,hum_ (Spearman’s ρ = −0.03, p = 0.82). In contrast to the initial sample, β_A,hum_ and β were also significantly associated (Spearman’s ρ = 0.64, p = 5 × 10^−7^), suggesting a more interdependent adjustment of learning and reputation mechanisms in this cohort. β_0,hum_ (Spearman’s ρ = 0.42, p = 0.002) correlated with earnings, but neither β_A,hum_ (Spearman’s ρ = −0.26, p = 0.06) nor β_U,hum_ (Spearman’s ρ = −0.14, p =0.3) did so significantly.

### Human-only games

Finally, we tested whether the computational architecture generalized to a simplified Human-only version of the task used for the subsequent TMS experiment. Across behavioral and fMRI samples (n = 77), the model combining U- and A-strategies again provided the best fit and predictive performance (Supplementary Figure 4), with significant contributions of both feedback-based updating (β_U_ = 0.26, 95% HDI [0.10, 0.42], p_MCMC_ = 0.0007) and reputation-based adjustment (βA = 0.19, 95% HDI [0.11, 0.27], p_MCMC_ < 0.0001). Thus, the underlying computational architecture generalized across independent datasets, including Human-only interactions.

### Oscillatory activity

Next, we examined the neurobiological features underlying the implementation of the identified negotiation strategies during instructed Human interactions. To isolate neural signatures associated with each strategy, we analyzed oscillatory brain activity using a model-based power modulation approach ^24–26^. In our previous work using the same task, we observed that alpha-band modulation in the right temporoparietal regions was linked to the anticipation of the partner’s response ^14,15,21,22^. Accordingly, the primary EEG analysis focused *a priori* on right temporoparietal electrodes. Using the trial-by-trial estimates derived from the model (for U strategy: β_U_ U_(t)_ and for A strategy: β_A_ A_(t)_), we quantified single-trial variations in oscillatory power. Because the U-strategy reflects feedback-based learning about the responder, we expected its neural modulation to emerge during the interval preceding the responder’s subsequent feedback. We found that trial-by-trial modulation in the U-strategy variable was associated with a positive modulation of alpha- and beta-band power during partner anticipation, extending from the decision period through the interval preceding the partner’s response, irrespective of game type. No such modulation was found for the A strategy (Figure 3 B).

**Figure 3.**
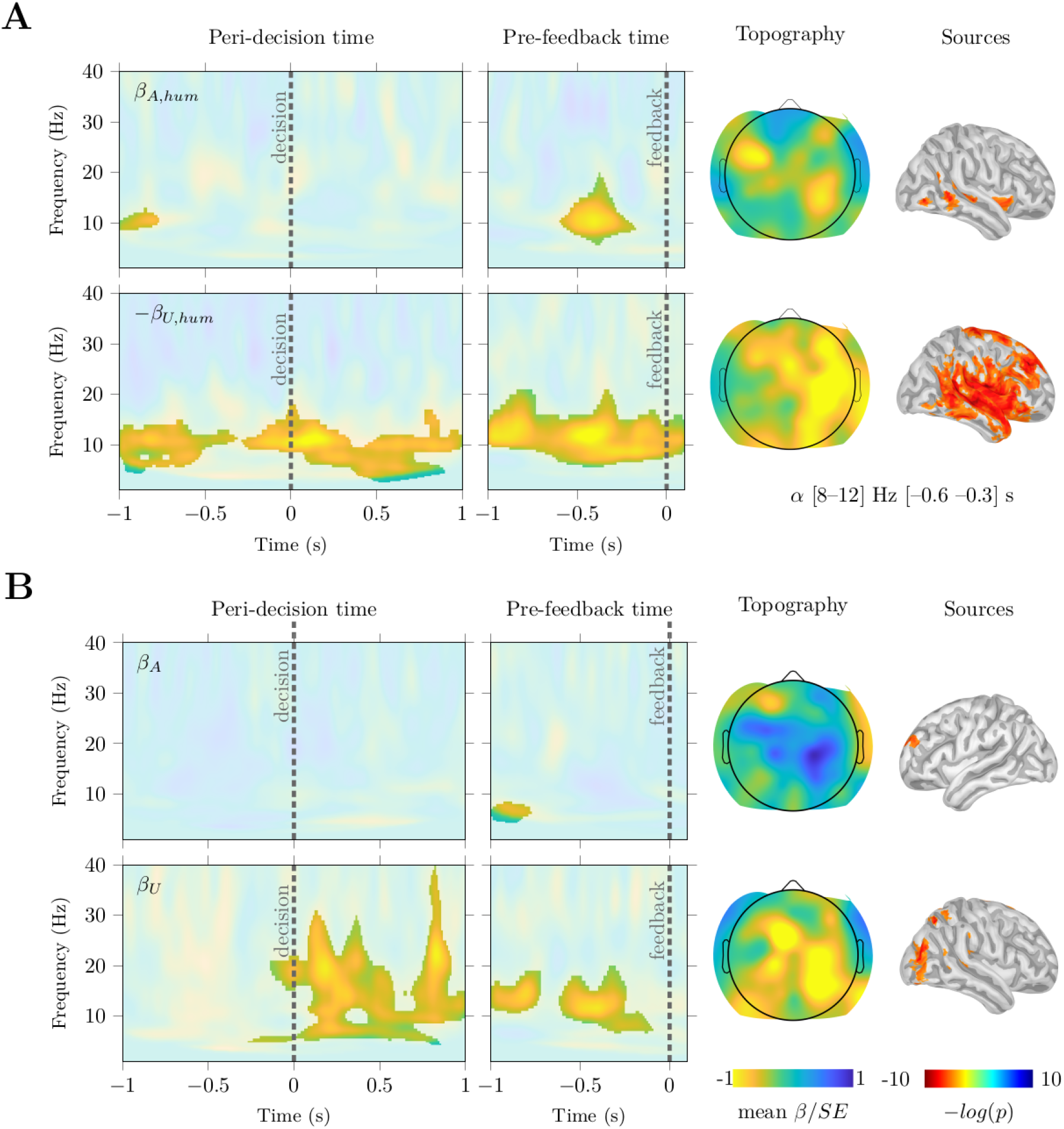
Alpha-band modulation of negotiation strategies in right temporoparietal electrodes. Time–frequency maps show oscillatory power modulation associated with trial-by-trial model-derived strategy estimates. Colors represent the mean normalized beta values across subjects (β/SE). Highlighted regions indicate clusters surviving a Wilcoxon signed-rank test against zero, followed by cluster-based permutation correction (p<0.001). Vertical dashed lines indicate decision (peri-decision panels) and partner feedback (pre-feedback panels). Topographic maps show the spatial distribution of alpha-band effects. Source estimates are displayed for regions surviving FDR correction (q < 0.05). **A**. Strategy modulation during instructed Human interactions, estimated from the interaction terms β_A,hum_ A_(t)_ H_(t)_ and −β_U,hum_ U_(t)_ H_(t)_. Both strategies show alpha-band modulation in right temporoparietal electrodes preceding partner feedback, with a stronger and more sustained effect for the U strategy extending from the decision period to the anticipation of the partner’s response. The A strategy shows a more localized modulation immediately preceding feedback. **B**.Strategy effects across games, estimated from β_A_ A_t_ and β_U_ U_t_. The U strategy is associated with increased alpha–beta power during partner anticipation, spanning the peri-decision and pre-feedback intervals. No significant modulation is observed for the A strategy. Topographies correspond to the alpha-band effect shown for the indicated time window.

We next explored the interaction between strategy engagement and human games, and found an extended alpha-band modulation associated with both strategies. Using the trial-by-trial estimates derived from the model (for the U-strategy: –β_U,hum_U_(t)_H_(t)_ and for the A-strategy: β_A,hum_ A_(t)_ H_(t)_), we found that strategic adaptation during human interactions was specifically linked to alpha power in this region. Interestingly, for the U strategy, despite the expected negative association with human engagement at behavioral levels, the model revealed a positive modulation in the alpha band related to this negative variation (Figure 3 A). Thus, stronger behavioral reductions in U-strategy engagement during Human games were associated with larger increases in alpha-band power. Hence, this pattern should not be interpreted as a reduction in cognitive processing underlying behavioral modulation. Rather, it indicates that the negative behavioral modulation of feedback-based updating was accompanied by increased alpha-band activity in right temporoparietal regions. Additionally, the A-strategy showed only a more localized modulation, restricted to the period immediately preceding the partner’s response. Source-level analyses revealed a strong alpha-band modulation associated with the U-strategy during human interactions, distributed across a network including the temporoparietal junction (TPJ), temporal cortex, and frontal regions. This spatial distribution overlaps with regions commonly implicated in mentalizing and social inference, encompassing TPJ, the superior temporal sulcus and inferior frontal gyrus ^27^.

### fMRI analysis

Then, using the human-only condition, we conducted an fMRI study to examine BOLD responses associated with the U- and A-strategies. Behaviorally, participants reproduced the expected decision patterns (see above, Human-only games section). Given the complex relationship between BOLD signals, neuronal activity^28,29^, and oscillatory dynamics^11,30,31^, we used multivariate Representational Similarity Analysis (RSA) to test whether local activity patterns reflected the computational structure of the model-derived strategies. For each participant, trials were sorted into 10 bins according to a common axis defined by the first principal component of the U- and A-strategy variables. We then computed representational dissimilarity matrices for each cortical parcel and tested whether these neural dissimilarity patterns were associated with the model-derived dissimilarity matrices for each strategy, while including reaction-time dissimilarity as a nuisance model accounting for differences in task demands.

This analysis revealed that the U-strategy was represented across several cortical parcels, including the right temporoparietal cortex (FDR-corrected, q < 0.05; Figure 4). This pattern was robust across variations in the number of bins and parcellation resolution (Figure 4 and Supplementary Figure 6). In contrast, no comparable representational pattern was observed for the A-strategy. As a functional validation of the fMRI dataset, we also examined the feedback period by relating BOLD activity to the reward obtained on each trial. Consistent with previous decision-making studies^32–34^, reward-related activity was observed in several regions, including the ventral striatum, ventromedial prefrontal cortex, and cerebellum (**Supplementary Figure 5**).

**Figure 4.**
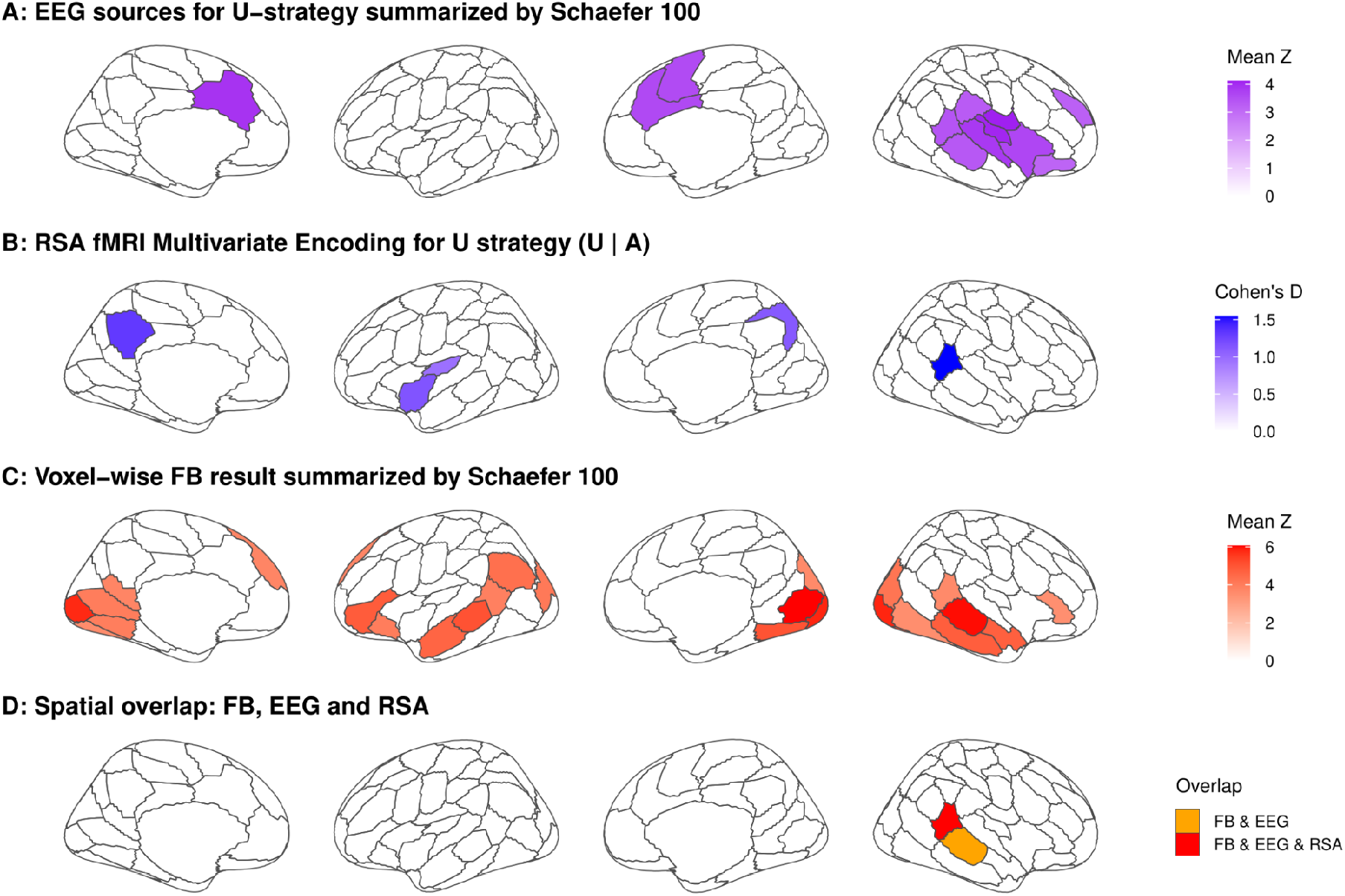
Multimodal convergence of U-strategy-related EEG and fMRI-RSA effects with regions engaged by a false-belief (FB) localizer. **A.** Source localization of alpha-band modulation associated with the U-strategy, projected onto the Schaefer 100-parcel atlas to facilitate visualization and comparison across modalities. Results are FDR-corrected at q < 0.05. **B.** Multivariate representational similarity analysis (RSA) of U-strategy-related representational structure across parcels of the Schaefer 100 atlas. Color represents Cohen’s d for parcels surviving FDR correction (q < 0.05). **C.** False-belief (FB) localizer activation for the contrast false belief > false photo. Voxel-wise results were projected onto the Schaefer 100-parcel cortical atlas to facilitate cross-modal comparison. Results are family-wise error cluster-corrected at p < 0.05. For panels A and C, color represents the mean z-statistic within each parcel. **D.** Spatial overlap across the three analyses, highlighting parcels jointly implicated by the false-belief localizer, U-strategy-related alpha source localization, and U-strategy fMRI-RSA.

**Figure 5.**
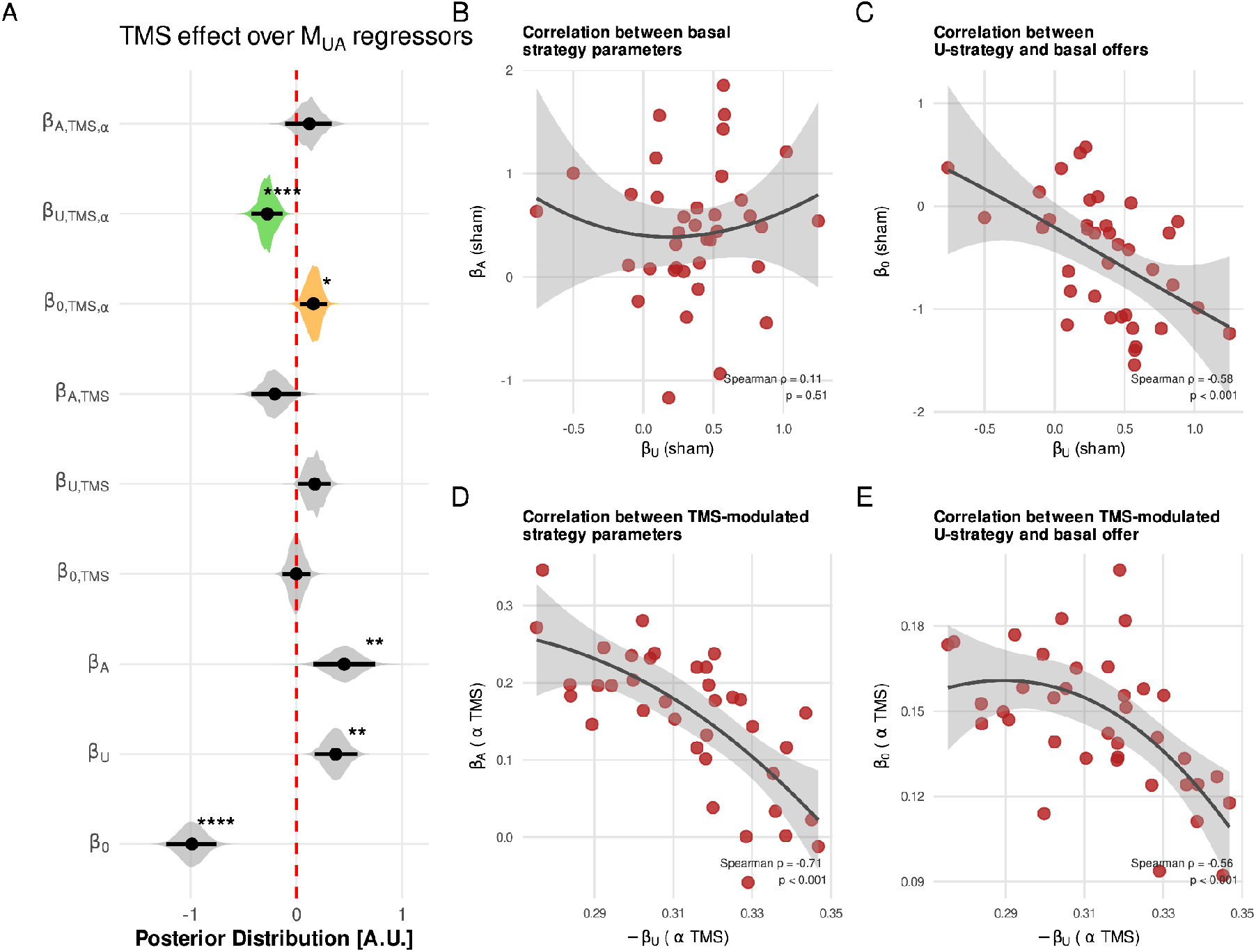
Rhythmic alpha-TMS selectively modulates feedback-based updating during social bargaining. **A.** Posterior distributions of regression coefficients for the computational strategy parameters and their modulation by stimulation. The TMS regressor represents the effect of active arrhythmic TMS relative to sham, thereby capturing non-frequency-specific effects of active stimulation. The TMSα regressor represents the additional effect of rhythmic alpha-TMS relative to active arrhythmic TMS, thereby isolating the alpha-specific stimulation effect. Rhythmic alpha-TMS reduced the contribution of the U-strategy relative to the active arrhythmic control (β_U,TMS,α_ p_MCMC_ <0.0001) and increased baseline offers (β_0,TMS,α_ p_MCMC_ =0.024). **B.** Relationship between U- and A-strategy parameters in the sham condition. **C.** Relationship between U-strategy engagement and baseline offers in the sham condition. **D.** Relationship between the alpha-specific modulation of the U- and A-strategy parameters, where both quantities correspond to the rhythmic alpha-TMS versus active arrhythmic TMS contrast. **E.** Relationship between the alpha-specific modulation of U-strategy engagement and baseline offers, again corresponding to the rhythmic alpha-TMS versus active arrhythmic TMS contrast. Points represent individual participants and correspond to posterior medians. Lines show LOESS fits for visualization. Spearman correlation statistics are reported in each panel. Asterisks indicate statistical evidence (*p<0.05, **p<0.01, ***p<0.001, ****p<0.0001).

### fMRI-guided rhythmic TMS modulation

To test whether perturbing temporoparietal alpha-band dynamics alters feedback-based updating during social negotiation, we designed a TMS experiment. First, participants underwent an fMRI session in which we identified the relevant anatomical regions to guide subsequent TMS targeting. To functionally localize each participant’s mentalizing network, we employed a well-established false-belief versus false-photograph task, which was first validated in an independent sample of 17 participants. Figure 4 illustrates the resulting activity in this sample, together with data from participants who subsequently underwent the TMS session. At the group level, the task showed activation in classical mentalizing network regions, including the temporoparietal junction (TPJ) and superior temporal sulcus in both hemispheres (Figure 4C). Since both EEG and fMRI results from the Ultimatum Game (UG) converged on the involvement of the right TPJ (Figure 4D), we targeted TMS to this region, including the angular gyrus and posterior superior temporal sulcus, using each participant’s individual peak of activity in the false-belief task within the group-level mask. We applied rhythmic TMS at an alpha frequency (five pulses delivered at 10 Hz, 100-ms inter-pulse interval, see Figure 1A) after the participant’s decision, but before they received their partner’s response, using arrhythmic and sham stimulations as controls. This timing was strategically chosen to avoid biasing the decision-making process itself while specifically modulating expectation formation regarding the partner’s likely response (we have used this strategy in other decision-making tasks, finding modulation in subsequent decisions^33^). By targeting this post-decision, pre-feedback window, we aimed to modulate the neural processes through which participants form predictions about their partner’s preferences, a critical component of the U-strategic learning that guides behavior, as indicated by our behavioral modeling in the Ultimatum Game.

In this experiment, 37 participants performed the Human-only version of the UG under three stimulation conditions: sham TMS, active arrhythmic TMS, and active rhythmic alpha-TMS. The two active conditions delivered the same number of pulses over the same stimulation period, with the arrhythmic condition serving as an active control for non-frequency-specific effects of TMS. In all behavioral and EEG models, stimulation was represented by two regressors. The TMS regressor was coded as 1 for both active conditions and 0 for sham, whereas the TMSα regressor was coded as 1 only for rhythmic alpha-TMS and 0 otherwise. With this coding, the TMS coefficient estimates the effect of active arrhythmic stimulation relative to sham, whereas the TMSα coefficient estimates the additional frequency-specific effect of rhythmic alpha-TMS relative to the active arrhythmic control.

Behaviorally, we first conducted a descriptive analysis correlating the programmed acceptance probability with the round number of each game. Consistent with previous findings, participants progressively adjusted their offers across rounds (β_round_ median = 0.003, HDI_95%_ [0.001, 0.006], p_MCMC_ = 0.009). This effect was not influenced by arrhythmic TMS stimulation (β_r,TMS_ median = 0.0006, HDI_95%_ [−0.001, 0.003], p_MCMC_ = 0.6), and rhythmic alpha-TMS stimulation ( β_r,TMSα_ = −0.001, HDI_95%_ [−0.0045, 0.0007], p_MCMC_ = 0.1). Interestingly, TMSα increased the baseline offers, as reflected by the modulation of the intercept term (β_0,TMSα_ median = 0.0675, HDI_95%_ [0.0174, 0.1163], p_MCMC_ = 0.006), an effect that was absent under arrhythmic TMS stimulation (β_0,TMS_ median = −0.0193, HDI_95%_ [−0.0636, 0.0230], p_MCMC_ = 0.3).

The effect observed in the first offers may indicate that TMSα modulated the U-strategy. To test this, we evaluated hierarchical models that included both strategic components. As expected, the model incorporating both strategies provided a better fit and predictive performance for the behavioral data (**Supplementary Figure 7**). The U-strategy showed a significant effect on participants’ behavior (β_U_ = 0.37, HDI_95%_ [0.17, 0.57], p_MCMC_ = 0.0011), as well as the A strategy (β_A_ = 0.45, HDI_95%_ [0.155, 0.744], p_MCMC_ = 0.004). Interestingly, in this model, TMSα significantly affected the U strategy, increasing the intercept term (β_0,TMSα_= 0.15, HDI_95%_ [0.02, 0.28], p_MCMC_ = 0.024) while decreasing the slope of the U-related parameter (β_U,TMSα_ = −0.27, HDI_95%_ [−0.43, −0.13], p_MCMC_ < 0.0001). In contrast, αTMS did not significantly influence the A strategy (β_A,TMSα_ = 0.12, HDI_95%_ [−0.10, 0.32], p_MCMC_ = 0.28).

To further characterize stimulation effects, we examined relationships among model parameters across participants. U- and A-strategy engagement were unrelated under sham TMS (Spearman’s ρ = 0.11, p = 0.51). For active arrhythmic TMS relative to sham, changes in U-strategy engagement covaried with both A-strategy modulation (ρ = 0.51, p = 0.0012) and baseline offers (ρ = 0.43, p = 0.0077). Similarly, for rhythmic alpha-TMS relative to active arrhythmic TMS, reduced U-strategy sensitivity was associated with changes in baseline offers (ρ = −0.56, p = 0.0003) and A-strategy modulation (ρ = −0.71, p = 1 × 10⁻⁶). Thus, alpha-TMS selectively modulated feedback-based updating while individual differences in this effect remained coupled to broader strategy changes.

### Rhythmic alpha-TMS modulates alpha oscillatory activity related to the U-strategy

We finally evaluated the effect of TMS on the EEG signal. To do so, we applied the same approach to that used in the EEG experiment, employing the latent variables for the U and A strategies to assess how the TMS interaction influenced their neural modulation trial-by-trial. Additionally, we included a TMS regressor to statistically rule out any direct stimulation effects that were not modulated by the behaviorally derived strategy variables. Although there is ongoing debate regarding the extent of local TMS effects, convergent evidence from both EEG and fMRI studies indicates that local responses tend to be more variable ^11,35^. In contrast, modulations in connected regions are more reliable ^36^, likely reflecting the propagation of physiologically meaningful perturbations along functional networks. In this context, and considering the need to minimize contamination from TMS-related artifacts, we examined right frontal electrodes, which were selected because they showed alpha-band modulation in the original EEG experiment and are plausibly connected to nodes of the mentalizing network.

Since the behavioral results showed that rhythmic alpha-TMS produced a pattern resembling that observed during Human games, we next examined the corresponding EEG effects using the same model-based approach, focusing on stimulation-related modulation of the U-strategy. In the sham condition, U-strategy-related activity showed alpha–beta modulation over frontal electrodes, consistent with the pattern observed in the original EEG experiment. No clear frequency-specific modulation was observed for the TMS regressor, which contrasts active arrhythmic TMS with sham and therefore captures non-frequency-specific effects of active stimulation. In contrast, the TMSα regressor, which contrasts rhythmic alpha-TMS with active arrhythmic TMS and therefore isolates the alpha-specific stimulation effect, revealed a robust alpha-band modulation over right frontal electrodes extending from approximately −700 ms to feedback onset. As an exploratory analysis, we examined right temporoparietal electrodes and observed a similar increase in alpha-band activity following rhythmic alpha-TMS (Supplementary Figure 8). Together, these findings indicate that the alpha-specific behavioral modulation of feedback-based updating was accompanied by increased alpha-band activity during anticipation of the partner’s response, consistent with the neural dynamics associated with the U-strategy.

### Rhythmic alpha-TMS modulation is associated with temporoparietal–frontal functional and structural connectivity

To assess whether the TMS effect was expressed at the network level, we performed exploratory functional and structural connectivity analyses. In the sham condition, alpha-band TPJ–frontal connectivity increased with U-strategy engagement (wPLI: β_U_ = 0.059, p = 0.006; Bayesian median β_U_ = 0.058, p_MCMC_ = 0.004). Rhythmic alpha-TMS significantly modulated this relationship relative to active arrhythmic stimulation (−β_U,TMSα_ = 0.060, p = 0.016; Bayesian median = 0.068, p_MCMC_ = 0.011), with the same pattern replicated using imaginary coherence (iCoh; β_U_: p = 0.025; −β_U,TMSα_: p = 0.031; Figure 7A).

**Figure 6.**
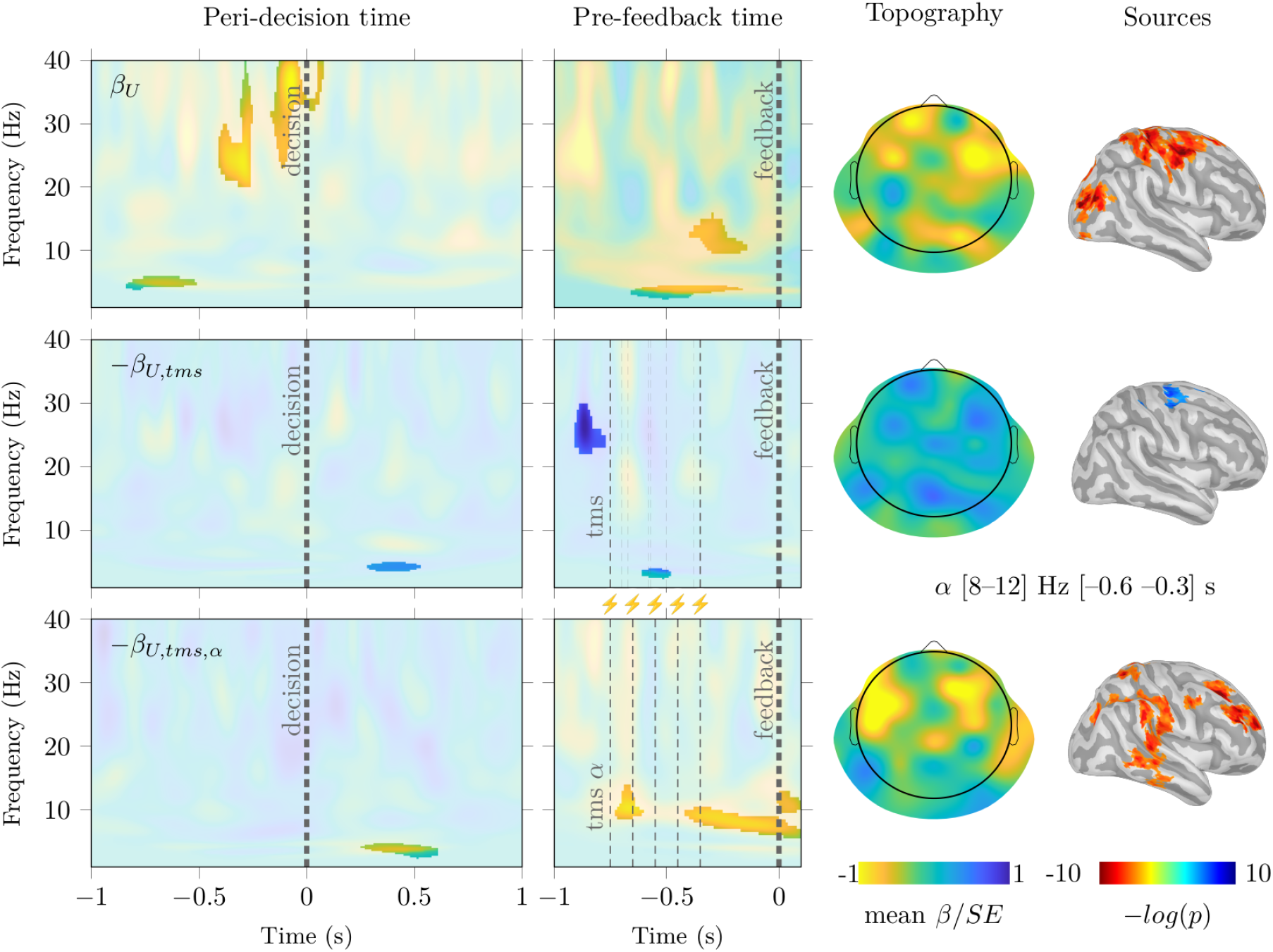
TMS modulates oscillatory activity related to U-strategy engagement over right frontal electrodes. Time–frequency maps show oscillatory power modulation associated with trial-by-trial model-derived U-strategy estimates and their modulation by TMS condition. Colors represent mean normalized beta values across participants (β/SE). Highlighted regions indicate clusters surviving a Wilcoxon signed-rank test against zero followed by cluster-based permutation correction (p < 0.001). Thick dashed vertical lines indicate the participant’s decision (peri-decision panels) and partner feedback onset (pre-feedback panels); thin dashed lines indicate TMS pulses. In the active arrhythmic condition, only the first and last pulses were fixed across trials, with intermediate pulses temporally jittered. Topographic maps show the scalp distribution of alpha-band effects, and source estimates show regions surviving FDR correction (q < 0.05). Upper panel, U-strategy-related modulation under sham TMS, showing frontal alpha–beta activity associated with feedback-based updating. Middle panel, active arrhythmic TMS modulation relative to sham, capturing non-frequency-specific stimulation effects. Lower panel, rhythmic alpha-TMS modulation relative to active arrhythmic stimulation, revealing a robust frontal alpha-band increase between the last TMS pulse and partner feedback.

**Figure 7.**
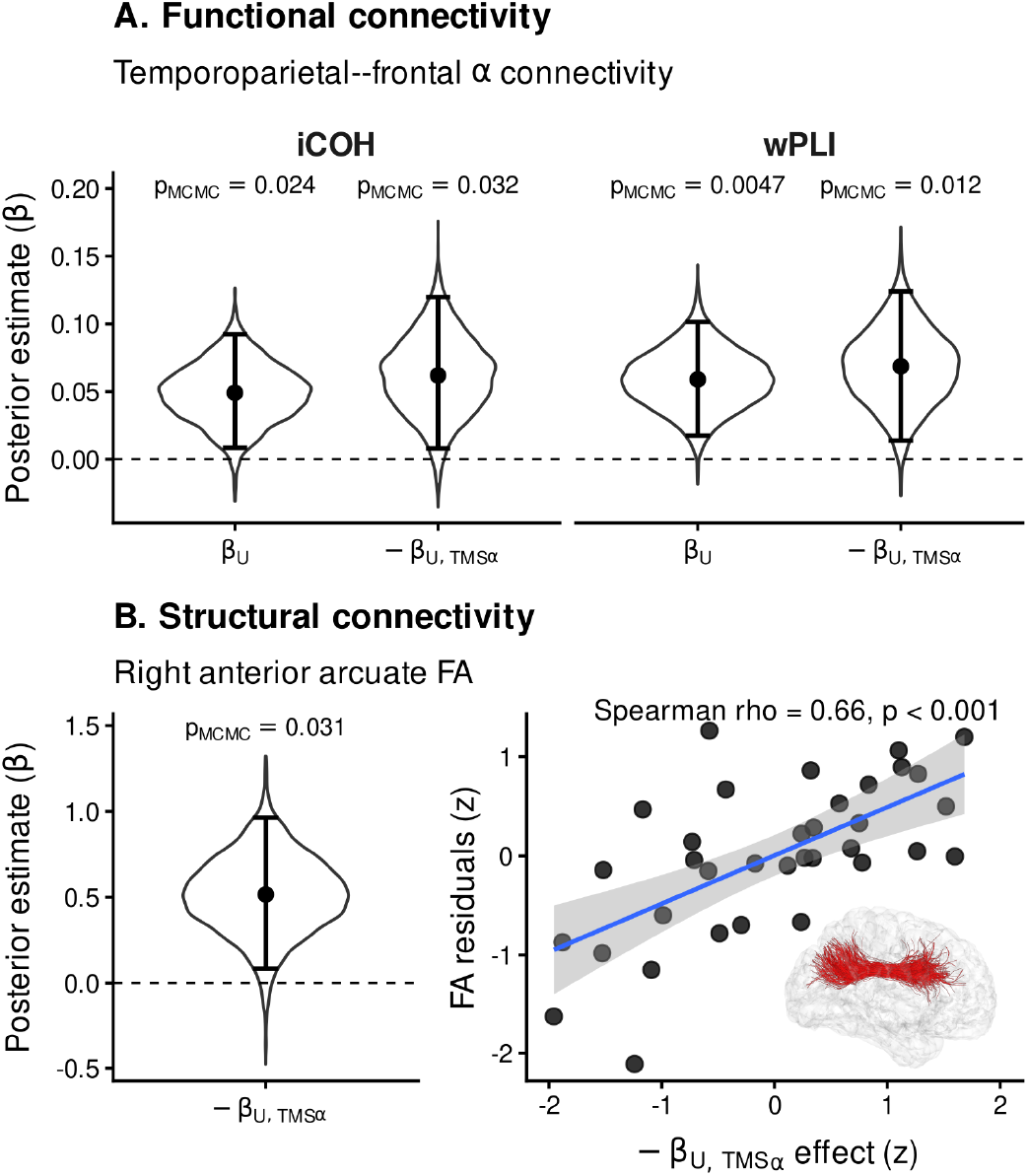
Functional and structural connectivity associated with alpha-TMS modulation of feedback-based updating. **A.** Posterior distributions of the effects linking U-strategy engagement to temporoparietal–frontal alpha-band connectivity, estimated using imaginary coherence (iCOH) and weighted phase-lag index (wPLI). β_U_ represents the association between U-strategy engagement and alpha-band TPJ–frontal connectivity, whereas −β_U,TMSα_ represents the alpha-specific modulation of this relationship for rhythmic alpha-TMS relative to active arrhythmic TMS. B. Structural connectivity analysis relating right anterior arcuate fasciculus fractional anisotropy (FA) to the alpha-specific behavioral modulation of feedback-based updating (−β_U,TMSα_), controlling for global FA and the remaining behavioral model parameters. The posterior distribution of the regression coefficient is shown on the left. The adjusted scatterplot on the right shows the relationship between −β_U,TMSα_ and right anterior arcuate FA residuals after covariate adjustment. The inset displays the right anterior arcuate tract. Shaded area indicates the 95% confidence interval. A–B, in violin plots, black dots indicate posterior medians, vertical bars indicate 95% HDIs, and dashed lines indicate zero. p_MCMC_ values are shown above each posterior distribution.

Structural connectivity analyses in 35 participants further showed that higher fractional anisotropy in the right anterior arcuate fasciculus was associated with stronger alpha-TMS modulation of the U-strategy (robust regression: −β_U,TMSα_ = 0.48, p = 0.005; Bayesian median = 0.51, p_MCMC_ = 0.03; Figure 7B). Similar associations were observed in other right, but not left, arcuate segments (Supplementary Figure 9). Together, these exploratory findings suggest that alpha-specific modulation of feedback-based updating is associated with functional and structural connectivity within a distributed temporoparietal–frontal network.

Together, these results identify feedback-based updating as the computational component most consistently linked to TPJ-centered alpha dynamics and show that rhythmic alpha-TMS selectively alters its behavioral expression.

## Discussion

Adaptive social bargaining requires dynamically weighting incoming feedback against more stable behavioral tendencies. Across independent samples, we identify feedback-based updating as a separable computational component with convergent temporoparietal alpha signatures across EEG and fMRI. Rhythmic alpha-TMS over individually localized TPJ sites selectively reduced this component relative to an active arrhythmic control and was accompanied by increased alpha-band activity and temporoparietal–frontal connectivity. Behaviorally, instructed social context did not uniformly shift strategy use; instead, individuals showed reproducible differences in how baseline behavior and feedback sensitivity were reweighted. Together, these findings link a specific computational operation in adaptive social behavior to temporoparietal alpha dynamics and provide perturbational evidence for their functional contribution to feedback weighting.

A central finding is that the behavioral impact of social context was expressed primarily through individual differences rather than a uniform Human-versus-Computer shift. Population-level interaction coefficients were close to zero, yet preserving the true responder-identity structure improved predictive performance relative to permuted condition labels, and participant-specific changes showed a highly structured relationship between baseline offers and feedback-based updating. The latent-mixture analysis further indicated that this pattern was expressed strongly in a substantial proportion of participants and, importantly, was reproduced in an independent sample. These results illustrate how averaging across individuals can obscure systematic forms of social adaptation: the same instructed context can alter the relative weighting of behavioral strategies in different ways across individuals.

Our computational results indicate that negotiation behavior can be decomposed into at least two distinct adaptive strategies. One strategy reflects learning about the responder from the outcomes of previous interactions, whereas the other captures adjustment based on the accumulated history of one’s own behavior, consistent with reputation-building strategies that could influence the counterpart’s future responses. This distinction suggests that repeated bargaining involves both adapting to others’ behavior and maintaining behavioral patterns that may shape subsequent interaction. Feedback-based updating was also systematically related to baseline offer levels, suggesting that adaptation emerges from the interplay between trial-by-trial evidence and relatively stable behavioral tendencies or expectations^10,37^.

An important implication of this computational framework is that strategic behavior did not correspond to immediate payoff maximization. When individuals consider not only others’ responses but also how their own behavior may shape future interactions, they may tolerate short-term costs or adopt apparently suboptimal offers to influence the partner’s expectations^38,39^, consistent with reputation-building accounts of repeated strategic interaction^16,40,41^. Accordingly, variability in offers that might otherwise appear as noise or inconsistency may partly reflect strategic exploration of the partner’s response contingencies^16,40,42,43^. In this context, our results suggest that repeated bargaining is supported by the flexible weighting of feedback-based updating and reputation-based adjustment, with substantial individual differences in how these strategies are deployed.

A particularly consistent individual-level pattern was the relationship between feedback-based updating and baseline offer modulation. Participants who increased their baseline offers more strongly during Human games tended to rely less on feedback-based updating, suggesting that strategic adaptation involves coordinated changes in relatively stable offer policies and trial-by-trial sensitivity to the partner’s responses. Baseline offers may therefore partly reflect stable expectations or social preferences upon which subsequent feedback-based updating operates, although the present data do not distinguish among specific normative interpretations such as fairness or social acceptability.

Although the processes supporting the inference of others’ preferences and intentions—commonly referred to as mentalizing—have been extensively studied using behavioral and neurophysiological paradigms^23,44,45^, less is known about how these processes are expressed during extended social exchanges in which individuals must continuously adapt their behavior to another agent. This question is central to second-person approaches to social neuroscience, which emphasize that social cognition emerges through active engagement, reciprocal exchange, and continuous updating rather than passive observation alone^1,2^. Repeated strategic games provide a controlled and computationally tractable setting in which to examine these processes. Our findings extend these accounts by showing that adaptation to instructed social context is not expressed as a uniform shift across individuals, but through structured differences in how behavioral tendencies and feedback-based updating are weighted during repeated bargaining.

Among the computational processes identified behaviorally, feedback-based updating showed the clearest convergent neural signature. Trial-by-trial U-strategy engagement was associated with alpha–beta modulation over right temporoparietal regions, while fMRI-RSA revealed corresponding representational structure in temporoparietal cortex overlapping regions engaged by the false-belief (FB) localizer. The absence of comparable effects for the A-strategy suggests that these neural signatures were preferentially related to the use of feedback for updating subsequent behavior rather than to strategic engagement more generally. This finding fits within a broader literature showing that social cognition relies on multiple oscillatory mechanisms across fronto-temporo-parietal networks, including theta activity during perspective-taking and inequity processing, beta activity associated with prosocial dispositions, and gamma-band fronto-temporoparietal coupling during social decisions^19,20,24,46–48^. Within this broader architecture, our results suggest a more specific and causal role for the right TPJ alpha oscillatory dynamics in regulating the influence of social feedback on belief updating. Predictive-inference accounts propose that adaptive behavior depends not only on detecting discrepancies between expected and observed outcomes, but also on regulating how strongly new evidence modifies existing beliefs ^45,49^. During negotiation, this requires balancing information obtained from the partner’s responses against more stable expectations and behavioral policies^50^. Alpha oscillations may contribute to this balance by regulating the effective weight and propagation of incoming information, consistent with proposed roles of alpha in information gating and precision weighting^15,51,52^. Recent work further suggests that increased alpha amplitude does not simply reflect cortical inhibition, but can actively shape ongoing neural processing^53–55^. Accordingly, higher alpha activity may reduce the behavioral impact of partner feedback, whereas lower alpha states may favor stronger trial-by-trial updating. This interpretation is consistent with previous findings showing increased temporoparietal alpha synchronization during socially meaningful interactions^15^ and smaller subsequent behavioral adjustments following rejection^14^. Together, these observations suggest that TPJ alpha dynamics may contribute to regulating how strongly social feedback influences future decisions during repeated strategic interaction^14–16,21,22^.

The perturbation experiment further supports a functional contribution of alpha-band dynamics to feedback-based updating. Critically, rhythmic alpha-TMS was compared directly with active arrhythmic stimulation matched in pulse number and stimulation period, allowing the TMSα contrast to isolate the additional frequency-specific effect of rhythmic stimulation. Rhythmic alpha-TMS reduced U-strategy engagement and increased baseline offers relative to the active arrhythmic control, while the corresponding EEG contrast revealed increased alpha-band activity during anticipation of the partner’s response. These effects were also expressed at the network level, with alpha-band functional coupling between temporoparietal and frontal regions varying with feedback-based updating and its modulation by rhythmic alpha-TMS. Structural connectivity analyses further linked individual differences in the behavioral effect of stimulation to white-matter connectivity within this network. Although the connectivity analyses were exploratory, their convergence with the behavioral and electrophysiological results suggests that alpha-specific modulation of feedback-based updating may involve a distributed temporoparietal–frontal system, rather than being restricted to the stimulated TPJ site.

The proposed gating process may be supported by distributed cortico-cortical and thalamo–cortical circuits involved in the generation and propagation of alpha-band activity ^56,57^. Corticothalamic loops can regulate thalamic gain and cortical synchronization, providing a potential mechanism through which information transmission is adjusted according to task demands and uncertainty^56–58^. Consistent with this possibility, alpha oscillations related to decision uncertainty have been associated with the integrity of corticothalamic white-matter pathways^59^. More broadly, alpha dynamics may coordinate top-down frontal influences with posterior temporoparietal processing, thereby regulating how strongly incoming information propagates through the cortical network to influence subsequent updating^53^. Although our data do not directly test a thalamic contribution, this systems-level account provides a plausible framework for understanding how alpha-band activity could regulate the impact of social feedback across distributed networks.

Several limitations qualify these conclusions. First, the Human-versus-Computer manipulation concerned instructed responder identity rather than actual online interaction with another participant. Although most participants reported experiencing the interaction as realistic, the debriefing did not directly establish that they believed the stated identity of the Human responder; these effects should therefore be interpreted as effects of instructed social context. Second, the latent-mixture analysis was exploratory in the initial sample, although its principal pattern was reproduced in an independent cohort. Third, the strategy-related univariate fMRI analysis did not yield significant clusters under conservative whole-brain correction, and the anatomical conclusions therefore rely primarily on multivariate representational analyses and cross-modal convergence. Finally, the functional and structural connectivity analyses were exploratory and should be confirmed in independent samples.

In summary, our findings show that feedback-based updating during repeated social bargaining is shaped by both individual differences in computational strategy and temporoparietal alpha dynamics. Social context did not impose a uniform strategy across participants; instead, individuals differed systematically in how baseline behavior and feedback sensitivity were reweighted. Across EEG, fMRI-RSA and frequency-specific TMS, feedback-based updating converged on alpha-band dynamics within a temporoparietal–frontal network. These findings provide a framework for understanding adaptive social behavior as the interaction between heterogeneous computational strategies and neural mechanisms that regulate the impact of feedback on future decisions.

## Methods

### Participants

Data were collected from 219 participants across 296 experimental sessions. The sample comprised five subgroups: the EEG sample (n = 40; 17–26 years; 20 women), each of whom completed an MRI and an EEG session; the replication sample (n = 48; 11–30 years; 22 women); the human-only sample (n = 77; 18–35 years; 30 women), including 34 fMRI sessions and 43 behavioral sessions; the false-belief probe sample (n = 17; 20–30 years; 10 women); and the TMS sample (n = 37; 18–37 years; 20 women), who each completed an MRI and a TMS-EEG session. Adult participants provided written informed consent. For minors, written informed consent was obtained from a parent or legal guardian and assent from the participant. These consent processes and all procedures were conducted in compliance with Chilean national legislation, institutional guidelines, and the Declaration of Helsinki statement of ethical principles for medical research involving human participants. All experimental procedures were approved by the Ethics Committee of Clínica Alemana—Universidad del Desarrollo, Chile (2018-03, 2020-67 and 2025-08).

### Behavioral task and measures

Participants performed a repeated Ultimatum Game as proposers. Each “game” comprised 20 consecutive trials with a fixed, anonymous responder. Across eight games, participants completed two conditions (except in the human-only sample): Human and Computer. In the Human condition, participants were told that they were playing against a human opponent online; in the Computer condition, participants were told that they were playing against a probabilistic algorithm. Importantly, in both conditions, opponents’ responses were generated by the same computational algorithm, allowing behavioral contingencies to be matched while manipulating instructed responder identity. The algorithm was implemented based on response patterns from human-to-human interactions documented in previous studies ^14^. Trials within each game were indexed as round (1–20).

After completing the task, participants underwent a brief debriefing interview about their experience of the Human and Computer conditions. Thirty-five of 40 participants (87.5%) spontaneously reported perceiving differences between the two conditions, most commonly describing one as more predictable, easier, or more difficult than the other. At the end of the interview, participants were also asked about the perceived realism of the online interaction. Only one participant reported that the interaction did not feel real, whereas the remaining participants reported experiencing the interaction as realistic.

### Derived behavioral index (primary outcome)

Because the algorithm provides a deterministic mapping from offers to acceptance probabilities P(acc|offer_(t)_), we used the z-normalized programmed acceptance probability of the offer on trial (t), p_(t)_ = z(P(acc|offer_(t)_)), as our primary continuous outcome. This measure captures adaptation more directly than raw offers by projecting them onto their behavioral consequence—the likelihood of acceptance—while supporting linear modeling with a continuous response and avoiding ceiling and floor effects associated with currency units.

### Computational predictors (latent strategies)

We formalized two complementary strategic mechanisms, updated within each game:

1. Feedback-history **updating** (U-strategy)

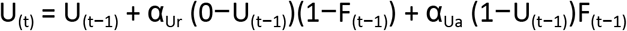

where F_(t−1)_ represents feedback on trial t−1 (0 = rejection, 1 = acceptance). α_Ur_ and α_Ua_ are learning rates for negative and positive feedback, respectively.

2. Reputation building **adjustment** (A-strategy)

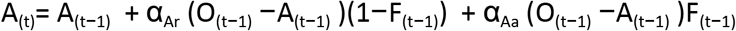

where O_(t−1)_ is the previous offer, and α_Ar_ and α_Aa_ are the reputation learning rates.

3. Alternative Reputation-Building Strategy (R strategy). As a control condition, we implemented a reputation-building model that did not distinguish between accepted and rejected offers. A key advantage of this formulation is that, when employed in conjunction with the U strategy, it completely disentangles the influence of the player’s own past actions (i.e., their previous offers) from the influence of other-regarding behavior (captured by the U strategy, which models responder responses).

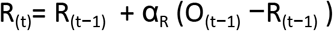

4. Latent-mixture control model. To assess whether the effect of opponent type was expressed homogeneously across participants or only in a subset of individuals, we implemented an additional hierarchical latent-mixture model. This model was designed as a control analysis to test whether the absence of a strong average Human-versus-Computer effect could reflect inter-individual variability in strategy modulation rather than the absence of a social-context effect. In this model, each participant was assigned a latent binary variable SC_s_ indicating whether that participant expressed a strategy shift between the Human and Computer conditions:

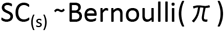

where π represents the population-level probability of showing a strategy shift. For participants with SC_(s)_=1, opponent type was allowed to modulate the computational parameters associated with the U-strategy, A-strategy, and baseline behavior. For participants with SC_(s)_=0, these Human-versus-Computer modulation terms were constrained to zero. Thus, the model simultaneously estimated both the proportion of participants showing strategy modulation and the direction and magnitude of the modulation. The behavioral response on each trial was modeled as a function of the computational predictors derived from the U- and A-strategies, together with their interaction with opponent type gated by SC_(s)_ (see below).

For the observation models, we modeled the acceptance probability as the outcome variable, using round and human game as regressors in a null model (M0).

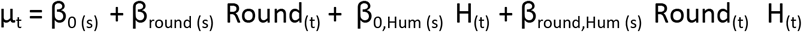

From the other models, each strategy has its own regressor. For the M_uah_ model, the equation is

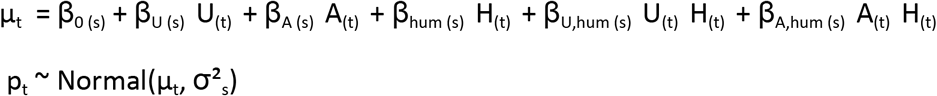

For the Latent-mixture control model, the equation is

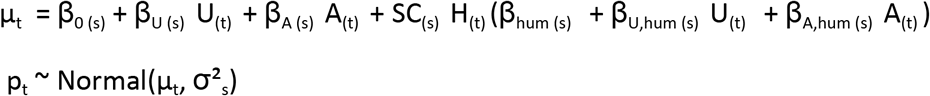

All beta parameters were modeled using a hierarchical structure with subject-specific coefficients drawn from group-level distributions, as follows:

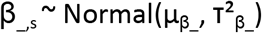

With the following prior

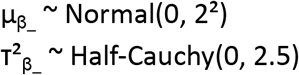

Regarding computational costs, it was not feasible to estimate all parameters in a hierarchical fashion. Therefore, to avoid biasing the model comparison, particularly for behavioral, EEG, and human-only experiments, only the learning parameters (α___) were estimated at the individual subject level. These were used to calculate the latent variables A, U, and R. In contrast, the β___ parameters were estimated hierarchically, allowing population-level inferences to be drawn exclusively from these parameters. We previously tested this approach in complex models with large samples ^11,32,59^.

### Bayesian inference and implementation

Models were implemented in JAGS (v4.1) called from R (v4.2) via the *rjags* and *runjags*. We used three chains, with 5000–10000 burn-in iterations and 1000 posterior samples per chain. Convergence was assessed by R-hat < 1.01, effective sample sizes, and trace diagnostics. We report medians, 95% HDIs, and posterior tail probabilities as an analog to p-value (p_MCMC_).

### Model comparison

Model fit was evaluated using Deviance Information Criterion (DIC) and Leave-One-Out Information Criterion LOOIC (package *loo* in R). Lower values indicate better fit and predictive accuracy.

### Earnings analyses

We summarized participants’ total earnings separately for the Human and Computer conditions and computed the difference score (Δ-earnings = Human − Computer) as an index of relative negotiation outcome. To examine whether individual variation in strategic modulation predicted performance, we modeled Δ-earnings as a function of the estimated strategy parameters (β_0,hum_ β_U,hum_, β_A,hum_). Specifically, we used Spearman rank correlations between individual strategy coefficients and Δ-earnings to capture monotonic relationships that are robust to non-normality.

### Anatomical data

For EEG, fMRI, and EEG-TMS experiments, all participants underwent high-resolution structural MRI scanning. Anatomical images were acquired on a 3T Siemens Skyra system (Siemens AG, Erlangen, Germany) equipped with a 45 mT/m gradient and a maximum slew rate of 200 mT/m/s. Each participant completed a T1-weighted MPRAGE acquisition (TR/TE = 2530/2.19 ms; TI = 1100 ms; flip angle = 7°; voxel size = 1×1×1 mm³), a T2-weighted sequence (TR/TE = 3200/412 ms; flip angle = 120°; echo train length = 258; voxel size = 1×1×1 mm³), and an axial 3D echo-planar imaging (EPI) (TR/TE = 8600/95 ms, 2 × 2 × 2 mm3 voxels, flip angle = 90°) with diffusion gradients applied in 30 non-collinear directions and two optimized b factors (b1 = 0 and b2 = 1000 s/mm2) with three repetitions. All imaging was acquired on a 3 T Siemens Skyra (Siemens AG, Erlangen, Germany) MR scanner with a gradient of 45 mT/m and a maximum slew rate of 200 mT/m/s. The T1 and T2 volumes consisted of 160 sagittal slices with isotropic resolution, providing full-brain coverage. T1/T2-corrected anatomical datasets were processed using the Human Connectome Project structural pipeline ^60^, yielding reconstructed scalp, cortical, and subcortical surfaces. Triangulated cortical meshes contained roughly 300,000 vertices per hemisphere before being down-sampled to ∼10,000 vertices for analysis. Additionally, a five-tissue segmentation was generated from the corrected T1- and T2-weighted images using the algorithms implemented in SimNIBS and SPM12.

### Functional MRI Data

During the fMRI session, functional data were acquired using a T2*-weighted echo-planar imaging sequence (TR/TE = 2390/35 ms; flip angle = 90°; voxel size = 4×3×3 mm) while participants performed either the UG or the False Belief task. For each participant, the resulting time series were nonlinearly normalized to the 2-mm isotropic MNI template using FNIRT from the FSL toolbox. To model neural activity during the False Belief task, two regressors were included: one for false-photograph trials and one for false-belief trials. The contrast between these two conditions was assessed to identify activity within the mentalizing network. Group-level statistical maps were generated using a mixed-effects approach (FSL FLAME1), and statistical inference employed cluster-based correction (z > 3.1 cluster-wise p < 0.05).

### fMRI representational similarity analysis

To identify cortical regions encoding the computational structure of the task, we performed a whole-cortex representational similarity analysis (RSA) using the Schaefer cortical parcellation. For each participant, trial-wise values of the U-strategy and A-strategy were extracted from the computational model and used to define a common representational axis. U- and A-strategy values were z-scored and submitted to a principal component analysis; the first principal component was then used to sort trials into 10 equally populated bins. For each bin, we built a separate event regressor using the trial onsets and durations convolved with a canonical hemodynamic response function. This produced one hemodynamic regressor per bin, plus an intercept term.

For each participant and each cortical parcel, we extracted the voxelwise BOLD time series from the preprocessed functional data without spatial smoothness. Within voxels of each parcel, we fit a voxelwise general linear model in which the binned hemodynamic regressors predicted the standardized BOLD signal. This yielded one multivoxel activation pattern per bin. We then computed a neural representational dissimilarity matrix for each parcel as one minus the correlation between multivoxel activation patterns across bins. Model representational dissimilarity matrices were constructed from the absolute pairwise differences between the mean U-strategy and A-strategy values across bins, while a reaction-time dissimilarity matrix was included as a nuisance regressor to control for differences in task demands. For each parcel, neural dissimilarity was modeled using a hierarchical Bayesian regression, with subject-specific intercepts and slopes and group-level parameters estimating the extent to which neural representational geometry was explained by U-strategy, A-strategy, and their interaction. All variables were standardized before model fitting. The main parameters of interest were the group-level effects of U-strategy and A-strategy dissimilarity on neural dissimilarity. Posterior distributions were estimated independently for each cortical parcel using three MCMC chains. For visualization and inference, we computed the posterior probability of each group-level effect having the opposite sign from its median direction and summarized effect size as the median group-level coefficient divided by the corresponding between-subject standard deviation. Parcels were retained when the posterior probability was below q=0.05 (FDR corrected). To assess robustness, we repeated the full analysis using 20 bins and a higher-resolution 200-parcel Schaefer atlas.

### DTI analysis

Data preprocessing and tractography followed the pipeline described in our previous work ^32,59,61–63^. Diffusion MRI data were processed using MRtrix3 ^64^. The preprocessing pipeline included image denoising, Gibbs ringing correction, motion and eddy-current distortion correction, and N4 bias field correction. The preprocessed diffusion images were then used to estimate the diffusion tensor model and extract fractional anisotropy (FA) maps. Specifically, fiber orientation distributions were estimated using 3-tissue response function estimation and multishell multi-tissue constrained spherical deconvolution ^65^. Whole-brain probabilistic tractography was performed using the iFOD2 algorithm with anatomically constrained tractography (ACT), using the following parameters: step size = 1 mm, maximum angle = 45°, minimum tract length = 25 mm, and maximum tract length = 250 mm. For each participant, 10 million streamlines were generated and subsequently filtered to 3 million streamlines using spherical-deconvolution informed filtering of tractograms (SIFT), reducing reconstruction biases and improving biological plausibility. T1-weighted and diffusion MRI images were co-registered using FSL. T1-weighted images were then nonlinearly normalized to MNI space, and individual tractograms were transformed accordingly. White-matter bundle segmentation was performed using an automated deep white matter segmentation algorithm based on Euclidean distances, together with the Deep White Matter (DWM) atlas, which defines 36 major white-matter bundles ^66^. The resulting streamline bundles were transformed back to each participant’s native space, where FA masks were computed for each bundle and participant to quantify inter-individual variability in tract microstructure.

### Structural connectivity analysis

To evaluate whether individual differences in white-matter microstructure were associated with the behavioral effects of alpha-rhythmic TMS, we fitted robust regression models using tract FA as the dependent variable. Tract microstructure was quantified using fractional anisotropy following our previous diffusion-MRI pipelines^32,62,63,67^. The primary predictor was the individual TMS-alpha modulation of the feedback-based learning strategy, −β_U,TMSα_, obtained from the hierarchical behavioral model. The primary tract of interest was the right anterior arcuate fasciculus, based on previous evidence linking the anterior arcuate/superior longitudinal fasciculus system to temporoparietal–frontal communication^68^. Additional exploratory analyses were performed for the remaining arcuate fasciculus segments to assess anatomical specificity. All continuous variables were standardized before model fitting. To account for potential confounding effects, the models included the remaining behavioral parameters from the computational model, including baseline offer parameters, TMS-related effects, and reputation-based learning parameters. Mean FA was also included as a covariate to control for global inter-individual differences in white-matter microstructure. To avoid circularity, mean FA was computed excluding the tract used as the dependent variable in each regression. Because these analyses were performed at the between-subject level and could therefore be sensitive to influential observations, we used robust regression models. Frequentist robust regressions were fitted using M-estimation, and Bayesian robust regressions were fitted using a Student’s *t* likelihood. Bayesian posterior distributions were summarized using the median, 95% highest density interval (HDI), and two-sided *p*_MCMC_ values. Associations were visualized using adjusted partial residual plots after accounting for all nuisance covariates.

### TMS-EEG

Stimulation targets were defined individually for each participant based on their BOLD activations during the False Belief task. For each participant, we identified the peak activation within the temporoparietal region, constrained to the population-level cluster derived from the independent sample of 17 participants who performed the same task. Stimulation sites were localized in native anatomical space using a neuronavigation system and each participant’s structural MRI. Individual coordinates were obtained by nonlinearly coregistering anatomical images to MNI space using FNIRT (FSL) and selecting the closest gray-matter voxel to the individually determined TPJ peak.

TMS coil position and orientation (yaw, pitch, and roll) were optimized such that the induced electric field was perpendicular to the cortical surface at the target, thereby maximizing effective current delivery ^69,70^. Stimulation intensity was set to the highest tolerable level for each participant within this region, corresponding to 100–120% of their individually measured motor threshold (45–80% of maximum stimulator output; mean = 60.6%, median = 60%). A 45-mm double coil (PMD45-EEG) was used to deliver pulses under three conditions: (1) an alpha-rhythmic condition (five pulses at 10 Hz, one every 100 ms), (2) an active arrhythmic condition (five pulses presented within a 400-ms window with jittered timing avoiding regular or 100 ms intervals between pulses), and (3) a sham condition. For both rhythmic and arrhythmic stimulation, the first and last pulses occurred at identical time points. Sham stimulation was delivered with the coil tilted, preventing effective cortical stimulation while preserving auditory and somatosensory artifacts.

EEG was recorded throughout both sessions using TMS-compatible equipment (BrainAmp DC 64-channel system, BrainProducts). Data were acquired from 64 scalp electrodes plus reference (FCz) and ground, using sintered Ag/AgCl TMS-compatible sensors. Signals were band-pass filtered from DC to 1000 Hz and digitized at 5000 Hz. Electrode impedance was kept below 5 kΩ and rechecked during task pauses to ensure stability. Electrode positions were estimated using the same neuronavigation system employed for TMS targeting.

### TMS electrophysiological analysis

EEG power was analyzed in two time windows: the interval surrounding participants’ decisions (when they submitted their offers) and the period surrounding the feedback corresponding to the partner’s response. All analyses were performed using the LAN toolbox for MATLAB (https://github.com/neurocics/LAN_current; https://matlab.mathworks.com/). Preprocessing was carried out in several stages. We first addressed the slow-decay component of the TMS artifact by extracting 3-second segments centered on TMS pulses, automatically identifying the interval from 10 ms before to 20 ms after each pulse peak, and removing this segment from the data. Independent Component Analysis (ICA) was then applied using the *runica* algorithm implemented in EEGLAB (https://sccn.ucsd.edu/eeglab/), allowing identification and removal of stereotyped components reflecting TMS-related artifacts and physiological activity such as blinks.

Next, the continuous EEG was segmented into the corresponding analysis windows. For each TMS pulse, the interval from −10 to 30 ms was excised and replaced using a shape-preserving piecewise cubic Hermite interpolant (pchip), ensuring smooth transitions without introducing oscillatory artifacts, and supplemented with Gaussian noise whose variance was estimated from a reference period (−55 to −15 ms before the first TMS pulse). This procedure effectively removed direct coil-induced artifacts and TMS-locked disturbances at nearby electrodes without generating discontinuities, a requirement for accurate subsequent time–frequency analysis ^69,71^.

Following artifact removal, signals were down-sampled to 1000 Hz and processed using a previously validated preprocessing pipeline ^15,21,24,33,48,72,73^. Continuous data were robustly detrended using a moving-median subtraction with a 2.5-second window (functionally equivalent to applying an ∼0.8 Hz high-pass filter but without introducing boundary artifacts). The data were then segmented, low-pass filtered at 45 Hz, and cleaned by removing ICA components corresponding to TMS-related artifacts, eye blinks, and eye movements. Artifact rejection was performed through a combination of automated and manual procedures. Automatic artifact detection was based on the following criteria: (1) voltage threshold exceeding ±150 μV or ±3 SD; (2) spectral threshold exceeding 2 SD in more than 10% of the 0.5–40 Hz frequency band; and (3) neighboring electrode correction falling below −3 SD. Trials identified as noisy were excluded. A subsequent manual inspection was conducted to verify and correct any remaining artifacts. Data from mastoid electrodes were excluded from all analyses.

Time-frequency decomposition was performed on each trial (t) and electrode (e) using complex Morlet wavelets (5-cycle width), applied to successive 10-ms time bins (tb) across the epoch spanning from −1.5 to 1.5 seconds relative to the decision and feedback events. For both time-windows of analysis, we conducted a first-level analysis using a general linear model over the power of each time-frequency bin in each EEG-TMS session. The linear regression includes the following regressors:

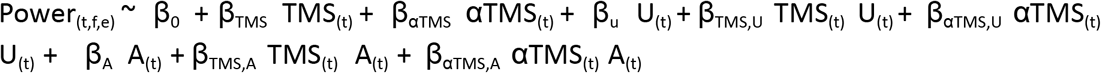

Then, for each session, the normalized beta value of each regressor (beta values / standard error) in each time-frequency bin was used in the second-level analysis. In this analysis, consistent modulation across subjects was tested in each bin using the Wilcoxon test. Then, the uncorrected significant differences were corrected for multiple comparisons using a cluster-based permutation test.

For the EEG experiments without TMS, the same analysis procedure was applied, with the exception that we used an EGI-amplified system with a 64-channel wet-net cap. The signals were recorded at 1000 Hz. The same preprocessing pipeline and segmentation were carried out. At the first level of analysis, we modeled the EEG signal using the following model:

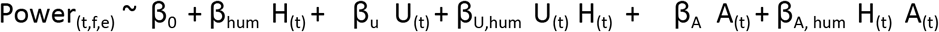

### TMS-EEG alpha-band connectivity

We performed an exploratory connectivity analysis between predefined temporoparietal and frontal electrode ROIs. The ROIs were defined using the same electrodes included in the time-frequency analyses. Connectivity was estimated at the single-trial level during the post-TMS/pre-feedback interval from −350 to 0 ms, corresponding to the period after the TMS burst and before the feedback. For each trial, EEG signals from temporoparietal and frontal electrodes were band-pass filtered in the alpha range. We then computed connectivity between all fronto-temporoparietal electrode pairs using two complementary phase-lag-based metrics: imaginary coherence (iCoh) and weighted phase-lag index (wPLI). These metrics were selected because they reduce the influence of zero-lag coupling and volume conduction relative to conventional coherence or phase-locking values. Connectivity values were then averaged across all fronto-temporoparietal electrode pairs to obtain one trial-wise alpha-band connectivity estimate per metric. The resulting trial-wise connectivity estimates were analyzed using mixed-effects models testing whether alpha-band fronto-temporoparietal coupling was modulated by the U-strategy regressor and by rhythmic alpha-TMS. The z-scaled Imaginary coherence and wPLI were analyzed separately to assess the robustness of the effect across connectivity metrics.

### Cluster-based Permutation Test

We employed a cluster-based permutation test to address multiple comparisons in our time-frequency analysis ^74^. Significant regions were identified by clustering neighboring sites showing the same effect (with a significance level of p < 0.05 in the statistical tests applied to either the time-frequency chart or the sources, such as the Wilcoxon test). Cluster-level statistics were computed as the sum of statistics from all sites within the cluster. We evaluated the significance of these clusters by comparing them to the largest cluster-level statistics in a permutation distribution. This distribution was generated by randomly permuting the original data. Specifically, we created null models for each participant, maintaining the original model structure while permuting the tested regressor. Following each permutation, we conducted the initial statistical test (e.g., Wilcoxon) and recorded the cluster-level statistics of the largest cluster from the permutation distribution. After 5,000 permutations, we estimated the cluster-level significance for each observed cluster as the proportion of values in the permutation distribution surpassing the cluster-level statistics of the corresponding cluster. For all time-frequency results, we highlighted all clusters that presented a p-corrected < 0.05. We indicated the highest p-value highlighted in each time-frequency chart to present more precise information.

### EEG Source Estimation

We estimated neural current density time series at each brain location using a minimum-norm estimated inverse solution algorithm (*LORETA*) with unconstrained dipole orientations. This estimation was performed individually for each trial, condition, and participant, employing Brainstorm software ^75^. We used a tessellated cortical mesh to model the brain based on each individual’s anatomy. This mesh was then used to estimate the distribution of current sources, with approximately 3x10000 sources positioned on the segmented cortical surface (three orthogonal sources at each spatial location). We employed a five-layer continuous Galerkin finite element conductivity model (FEM), as implemented in DUneuro software^76^, along with a physical forward model. We multiplied the recorded raw EEG time series from the electrodes by the inverse operator to obtain cortical activity estimates at the cortical sources. This operation yielded the estimated source current at the cortical surface as a function of time. Importantly, this transformation is linear and does not alter the frequencies of the underlying sources, enabling us to conduct time-frequency analyses directly in the source space. Within this source space, we simplified each vertex’s dipole to a single time series by applying a variance-weighted projection. For each vertex, we computed the variance of each of the three orthogonal dipole components across time and used the normalized variances as weights to obtain a weighted linear combination of the three time series. This method approximates the principal component analysis (PCA) but is computationally faster, as it does not require eigenvalue decomposition. Subsequently, we performed frequency decomposition using the wavelets transform. To ensure the reliability of our results, we only present source estimations if statistically significant differences are observed at both the electrode and source levels, thereby accounting for multiple comparison corrections.

## Data Availability

The complete minimal data set underlying the results used in our study is available in the public repository on GitHub (https://github.com/neurocics/Figueroa-Vargas2026). The additional toolbox and codes used in the analysis are available on our lab github site (https://github.com/neurocics/LAN_current).

## Acknowledgements

This work was supported by Grants 1251073 (PB), 11261678 (AF-V), 1261590 (PS-I), 1211227 (PB), 1181295 (PB), and 11140537 (PB) AC3E CIA250006 (PG), Beca Doctorado Nacional 2025-21251976 (SN) from the Fondo Nacional de Ciencia y Tecnología (FONDECYT) from Agencia Nacional de Investigación y Desarrollo (ANID). The funders had no role in study design, data collection, and analysis, decision to publish, or preparation of the manuscript.

## Supplementary Figures

**Supplementary Figure 1.**
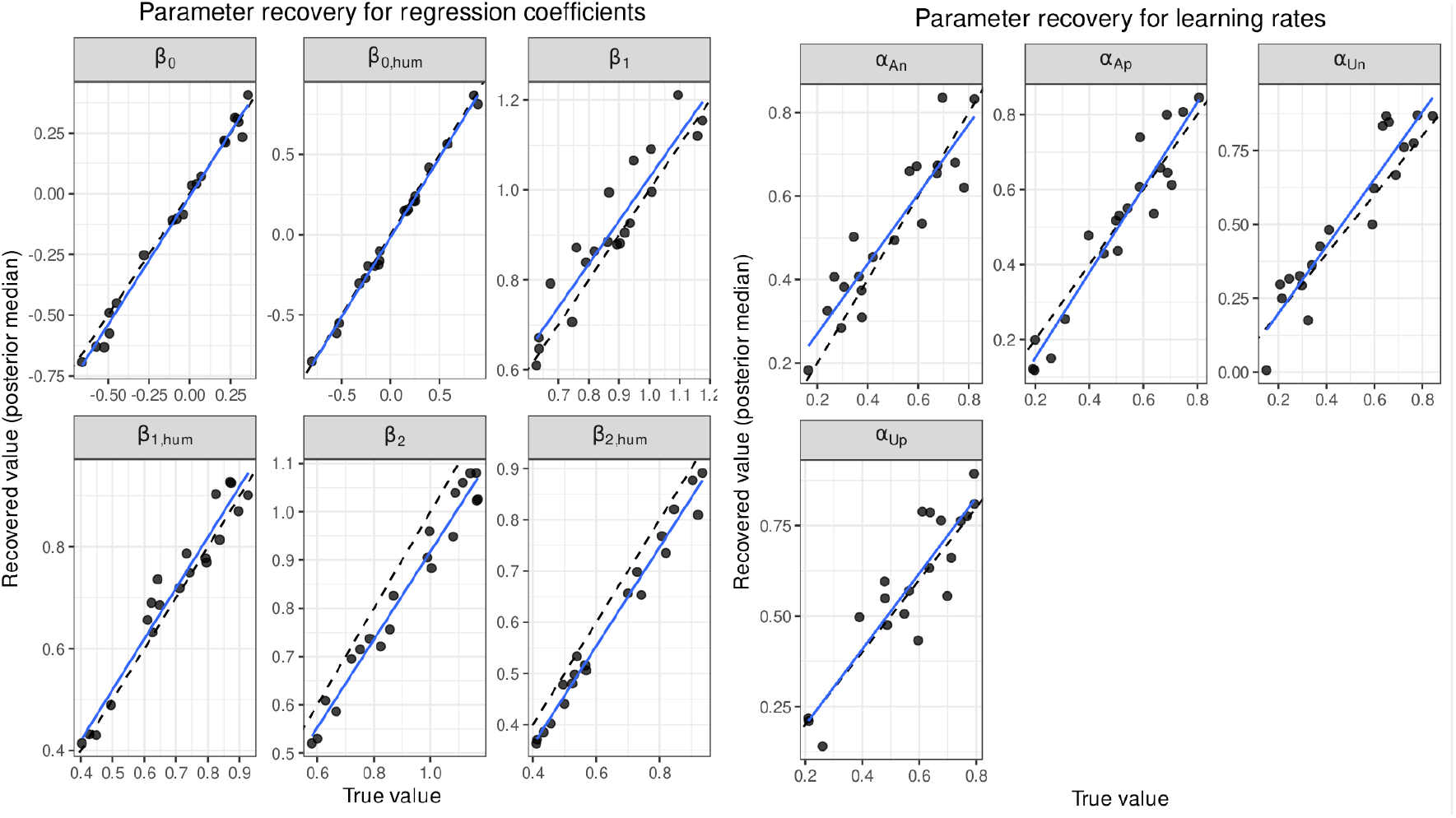
Parameter recovery analysis. Parameter recovery was assessed using simulated datasets generated from the full model. For regression coefficients, learning rates were fixed (α_Up_=α_Un_=α_Ap_=α_An_=0.5), and β parameters were sampled across a wide range of values. For learning rates, regression coefficients were fixed at non-zero values (β_1_, β_1,hum_, β_2_, β_2,hum_) to ensure sufficient influence of latent variables on observed behavior. Each panel shows the relationship between true generating values and recovered posterior medians across simulations. Dashed lines indicate identity (y=x). Overall, parameters were reliably recovered, with high correspondence between true and estimated values.

**Supplementary Figure 2.**
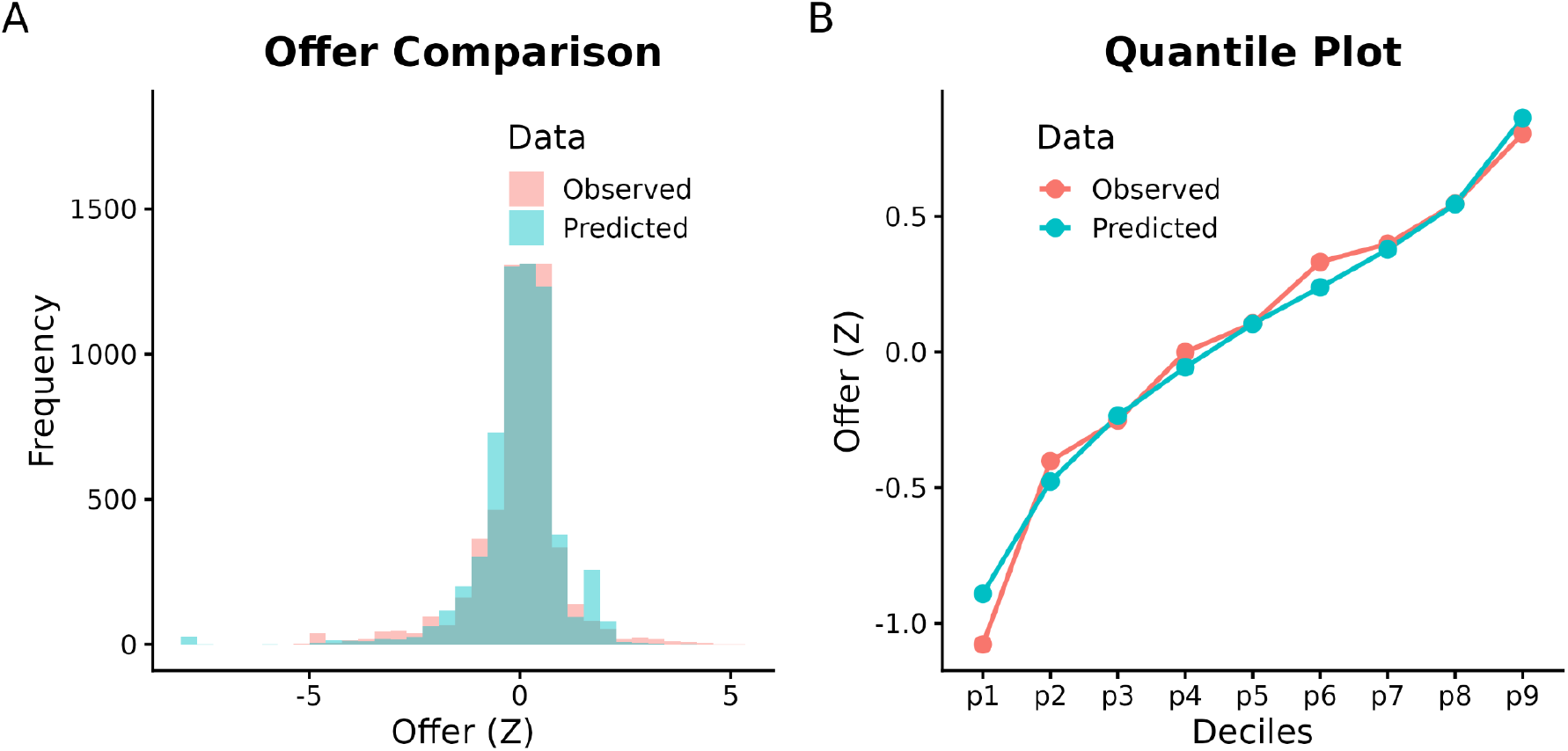
Model fit to offer distributions. (A) Histogram comparing observed and model-predicted offers (Z-scored) across trials. (B) Quantile–quantile plot showing observed and predicted offer distributions across deciles. The model accurately reproduces both the overall distribution and the quantile structure of the observed data, indicating a good fit at both central tendency and distributional levels.

**Supplementary Figure 3.**
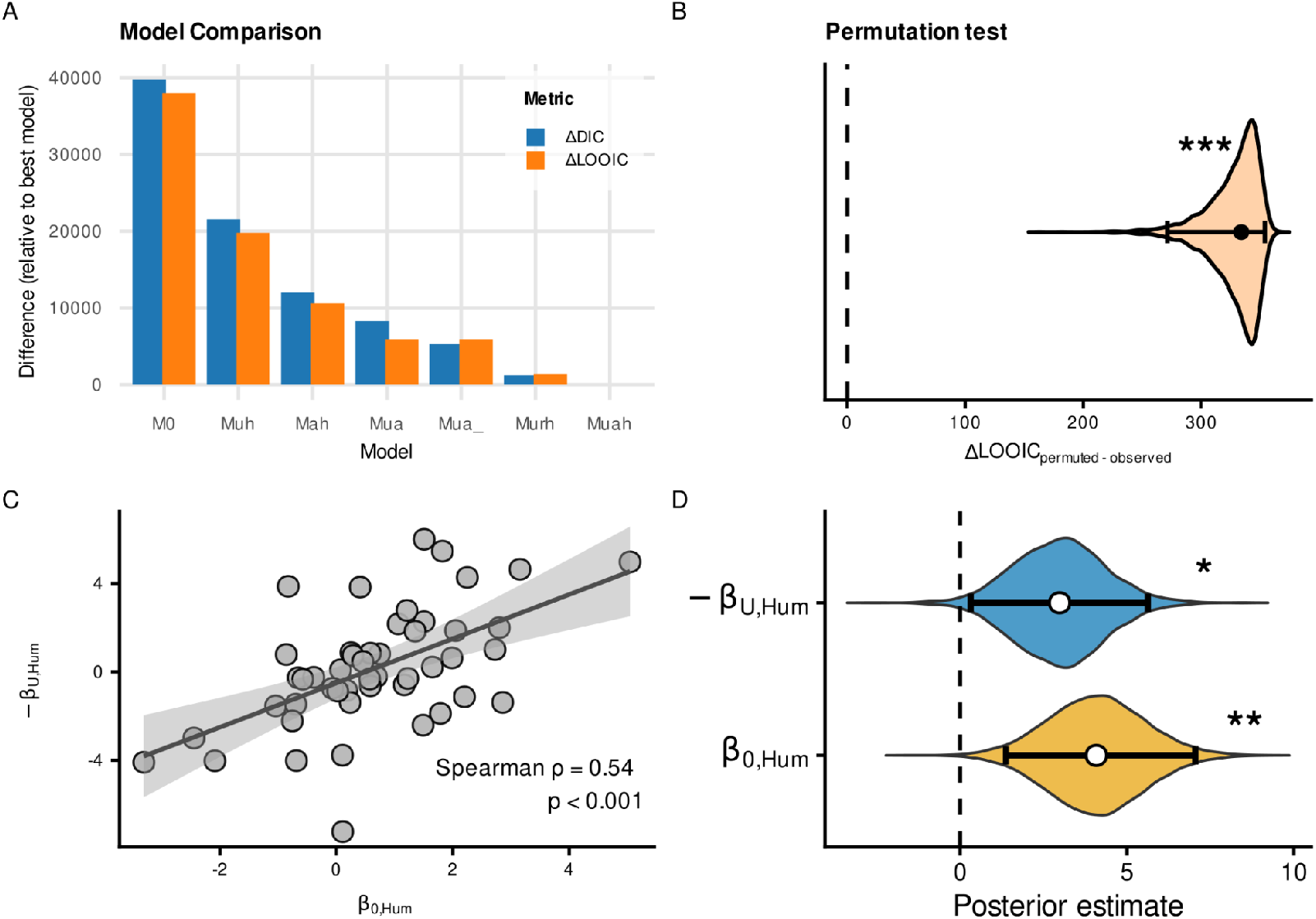
Behavioral strategies during social negotiation in a replication sample (n=48). A, Model comparison across candidate computational models. Bars show differences in model fit and predictive accuracy relative to the best-fitting model, quantified using ΔDIC and ΔLOOIC, respectively. The model combining feedback-based updating (U-strategy) and reputation-based adjustment (A-strategy), together with their modulation during instructed Human interactions, provided the best overall account of behavior. **B,** Permutation test assessing whether the predictive advantage associated with responder identity reflected meaningful Human-versus-Computer condition structure. The violin shows the distribution of ΔLOOIC values obtained after permuting condition labels across games within participants, expressed relative to the model fitted using the observed condition labels (permuted − observed). Positive values indicate poorer predictive performance after permutation. The dashed line indicates ΔLOOIC = 0. C, Individual differences in strategy modulation during Human games. Participants who increased their baseline offers more strongly during Human interactions (β_0,hum_) also showed a greater reduction in feedback-based updating, represented in the figure as _−βU,hum_. Points represent individual participants; the line and shaded area show the linear fit and 95% confidence interval for visualization. Spearman’s correlation is reported in the panel. D, Posterior distributions from the hierarchical Bayesian latent-mixture model for the Human-related increase in baseline offers (β_0,hum_) and reduction in feedback-based updating (β_U,hum_). White circles indicate posterior means and horizontal bars indicate 95% highest-density intervals (HDIs); the dashed vertical line indicates zero. Asterisks indicate posterior evidence against zero (*p_MCMC_<0.05, **p_MCMC_<0.01, ***p_MCMC_<0.001).

**Supplementary Figure 4.**
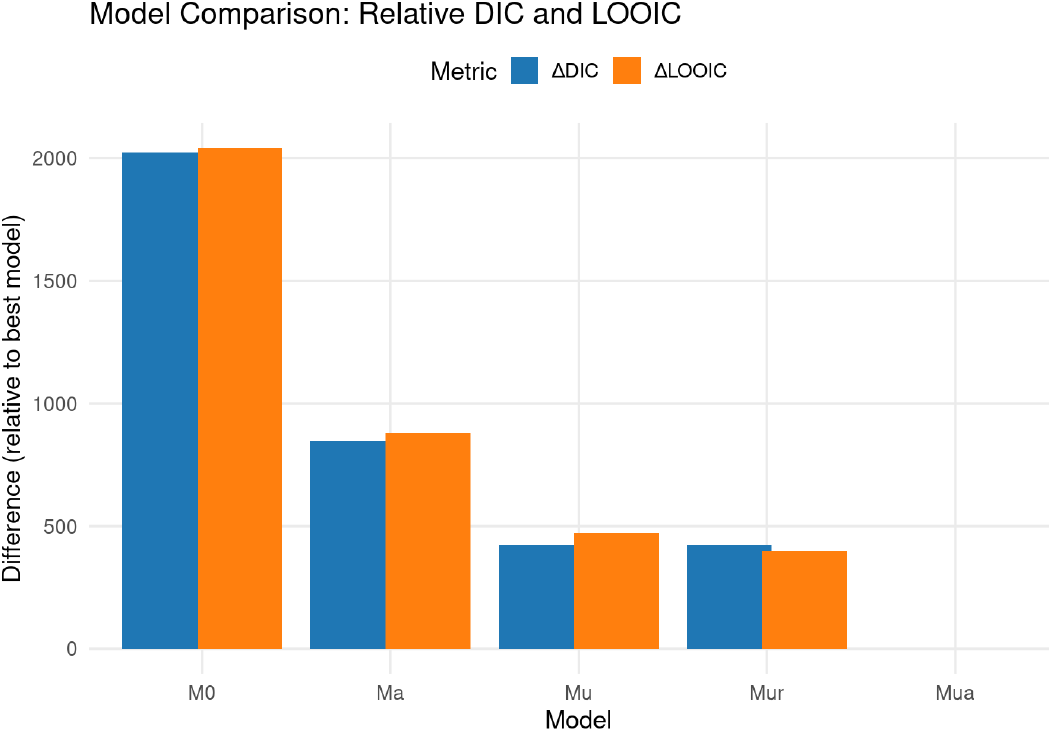
Behavioral strategies during social negotiation in a human-only sample (n=77). Model comparison across candidate computational models. Bars show differences in model fit relative to the best-fitting model quantified using ΔDIC and ΔLOOIC. The model that includes both feedback-based learning (U-strategy) and reputation-based updating (A-strategy) provides a better fit and better predictions.

**Supplementary Figure 5.**
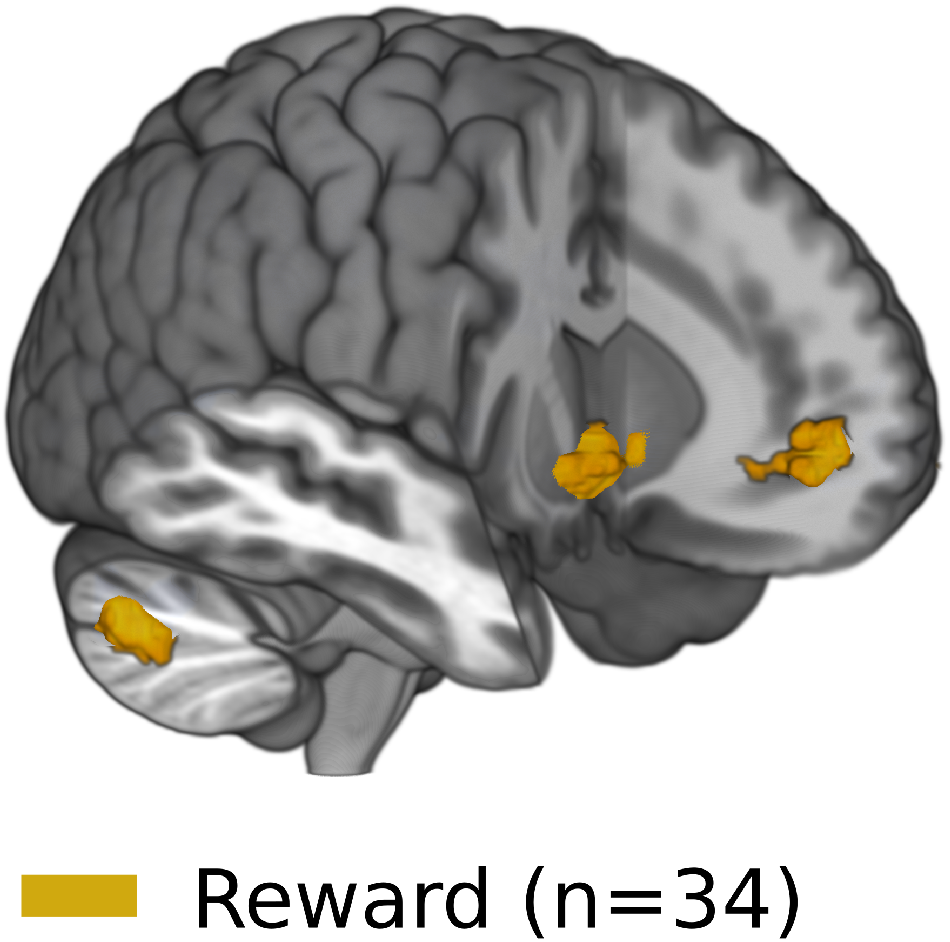
Univariate fMRI localization of reward-related activity. Whole-brain maps show BOLD responses associated with trial-by-trial reward during feedback, with significant effects primarily observed in the ventral striatum, ventromedial prefrontal cortex, and cerebellum. The first-level model was specified as BOLD ∼ 1 + reward + rejection + acceptance, with the reward regressor z-scored prior to model estimation. Subject-level reward effects were then entered into a second-level mixed-effects analysis. Statistical maps were thresholded using a cluster-forming threshold of z > 3.1 and corrected for multiple comparisons at the cluster level. All displayed clusters survived cluster-wise correction at p < 0.01.

**Supplementary Figure 6.**
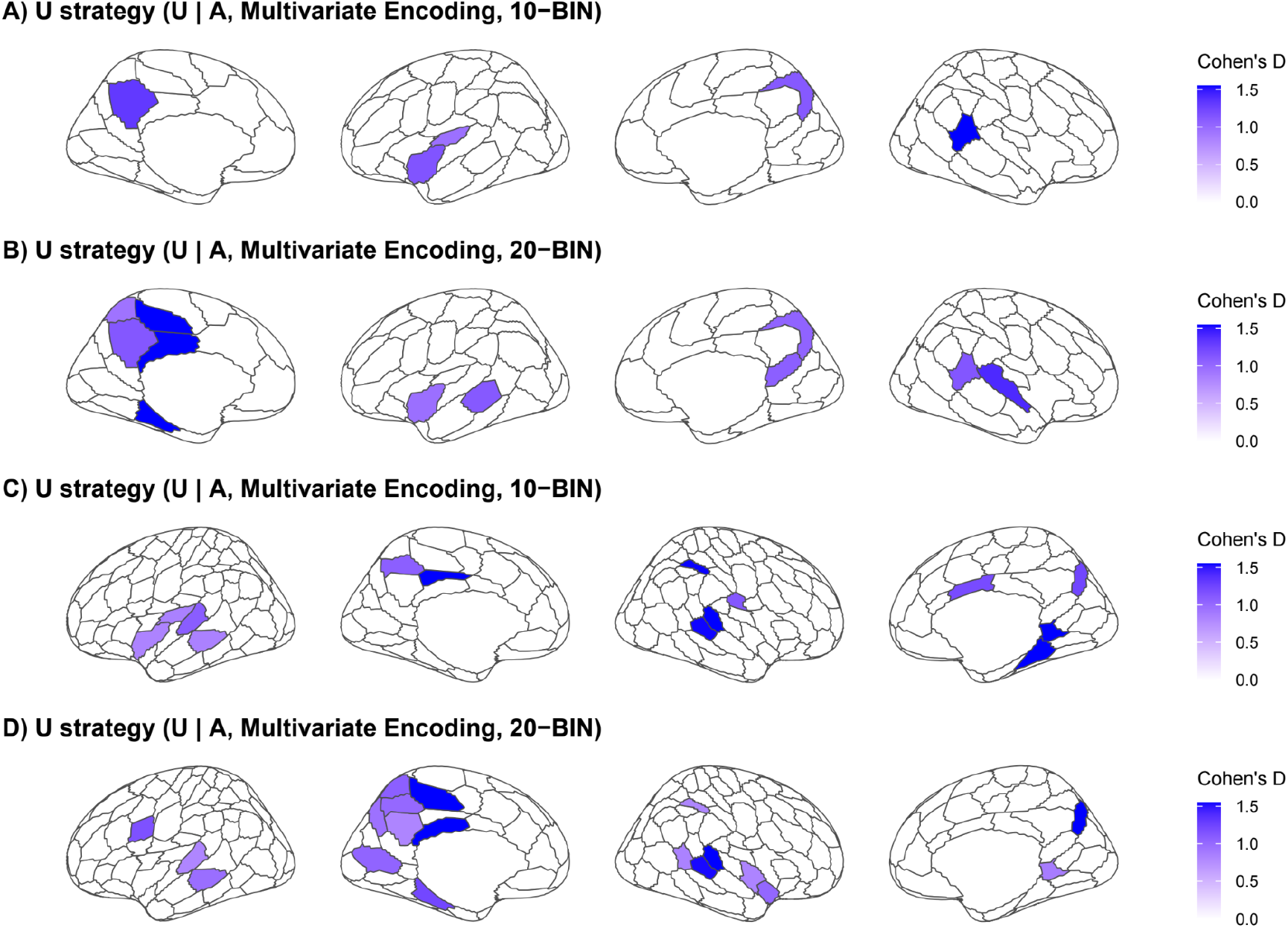
Robustness of fMRI-RSA encoding of the U-strategy across binning schemes and parcellation resolutions. Multivariate representational similarity analysis (RSA) was used to identify cortical parcels whose representational geometry encoded the feedback-based updating strategy, U, while controlling for the reputation-based A-strategy, the U × A interaction, and reaction-time dissimilarity. To assess robustness, the analysis was repeated using two temporal binning schemes, 10 and 20 bins, and two Schaefer cortical parcellation resolutions, 100 and 200 parcels. Panels show **A**, Schaefer 100 with 10 bins; **B**, Schaefer 100 with 20 bins; **C**, Schaefer 200 with 10 bins; and **D**, Schaefer 200 with 20 bins. Color indicates Cohen’s *d* effect size for significant parcels after FDR correction. The repeated involvement of temporoparietal regions across specifications supports the robustness of the anatomical localization of U-strategy encoding.

**Supplementary Figure 7.**
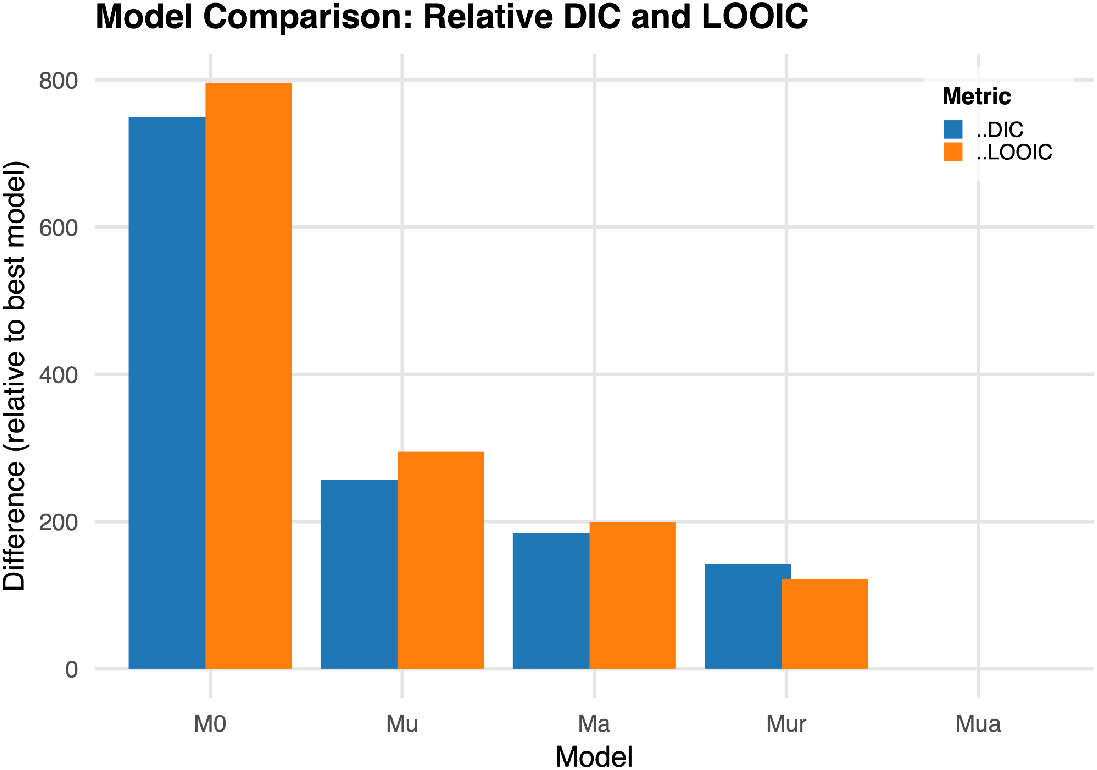
Behavioral strategies during social negotiation in a TMS sample (n=37). Model comparison across candidate computational models. Bars show differences in model fit relative to the best-fitting model quantified using ΔDIC and ΔLOOIC. The model that includes both feedback-based learning (U-strategy) and reputation-based updating (A-strategy) provides a better fit and better predictions.

**Supplementary Figure 8.**
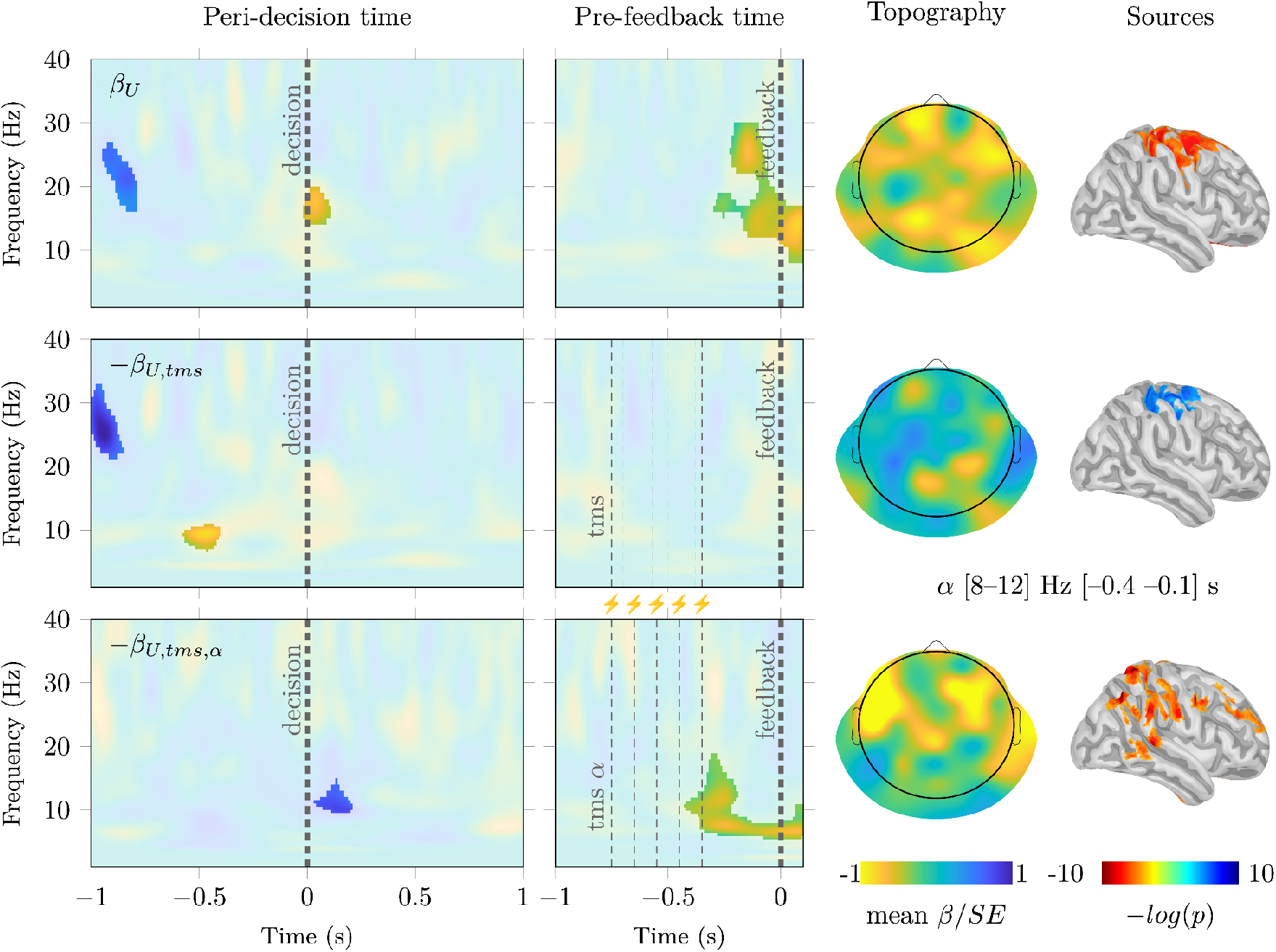
TMS modulates oscillatory activity related to U-strategy engagement over right temporoparietal electrodes. Time–frequency maps show oscillatory power modulation associated with trial-by-trial model-derived U-strategy estimates and their modulation by TMS condition. Colors represent mean normalized beta values across participants (β/SE). Highlighted regions indicate clusters surviving a Wilcoxon signed-rank test against zero followed by cluster-based permutation correction (p < 0.001). Thick dashed vertical lines indicate the participant’s decision (peri-decision panels) and partner feedback onset (pre-feedback panels); thin dashed lines indicate TMS pulses. In the active arrhythmic condition, only the first and last pulses were fixed across trials, with intermediate pulses temporally jittered. Topographic maps show the scalp distribution of alpha-band effects, and source estimates show regions surviving FDR correction (q < 0.05). Upper panel, U-strategy-related modulation under sham TMS, showing frontal alpha–beta activity associated with feedback-based updating. Middle panel, active arrhythmic TMS modulation relative to sham, capturing non-frequency-specific stimulation effects. Lower panel, rhythmic alpha-TMS modulation relative to active arrhythmic stimulation, revealing a robust frontal alpha-band increase between the last TMS pulse and partner feedback.

**Supplementary Figure 9.**
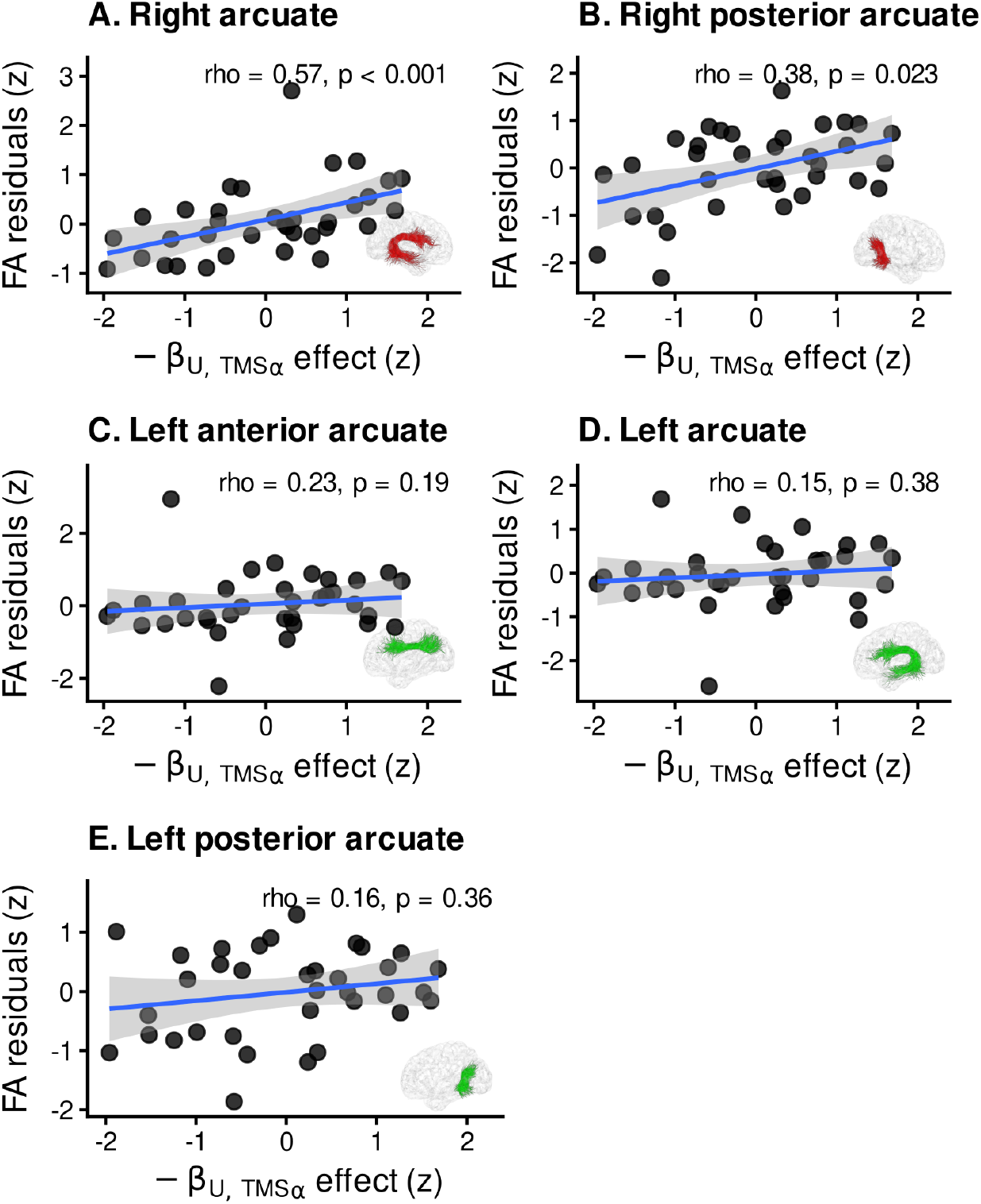
Specificity of the association between arcuate fasciculus FA and TMS-alpha modulation of feedback-based updating. Adjusted associations between the TMS-alpha modulation of the U-strategy, −β_U,TMSα_, and fractional anisotropy (FA) residuals in different segments of the arcuate fasciculus. **A.** Right arcuate fasciculus. **B.** Right posterior arcuate fasciculus. **C.** Left anterior arcuate fasciculus. **D.** Left arcuate fasciculus. **E.** Left posterior arcuate fasciculus. FA residuals were obtained after controlling for global FA and the remaining behavioral model parameters. Each point represents one participant. Blue lines show the fitted linear association, and shaded areas indicate 95% confidence intervals. Insets show the tract segment included in each analysis. Spearman’s ρ and corresponding *p*-values are shown in each panel.Spearman’s ρ values are shown for descriptive visualization; inferential statistics reported in the main text were obtained from the robust regression models described in Methods.

